# How many and which species to plant? A multi-trait-based approach to select species to restore ecosystem services

**DOI:** 10.1101/2020.10.29.359455

**Authors:** Paula Kiyomi Tsujii, André Ganem Coutinho, Anderson M. Medina, Nathan J. B. Kraft, Andres Gonzalez-Melo, Pedro Higuchi, Sandra Cristina Müller, Ülo Niinemets, Valério D. Pillar, Enio E. Sosinski, Marcos B. Carlucci

**Affiliations:** Programa de Pós-Graduação em Ecologia e Evolução, Universidade Federal de Goiás, Goiânia, GO, Brazil; Department of Ecology and Evolutionary Biology, University of California, Los Angeles, CA, USA; Facultad de Ciencias Naturales y Matemáticas, Universidad del Rosario, Bogotá, Colombia; Departamento de Engenharia Florestal, Universidade do Estado de Santa Catarina. Lages, SC, Brazil; Departamento de Ecologia, Universidade Federal do Rio Grande do Sul. Porto Alegre, RS, Brazil; Estonian University of Life Sciences, Kreutzwaldi 1, Estonia; Embrapa Recursos Genéticos e Biotecnologia, Brasília, DF, Brazil; Laboratório de Ecologia Funcional de Comunidades (LABEF), Departamento de Botânica, Setor de Ciências Biológicas, Universidade Federal do Paraná, Curitiba, PR, Brazil

**Keywords:** ecosystem function, ecosystem resilience, functional diversity, functional redundancy, functional restoration, functional trait, multifunctionality, species selection

## Abstract

It has been increasingly argued that ecological restoration should focus more on targeting ecosystem services than on species composition of reference ecosystems. In this sense, the role that species play on community assembly and functioning through their functional traits is very relevant, because effect traits mediate ecosystem processes, ultimately resulting in provision of ecosystem services. One major challenge in ecological restoration is to know which species to use that will deliver the target ecosystem services. We developed an algorithm to select the minimum set of species that maximize the functional richness (FRic) and the functional redundancy (FR) of the restored community, a proxy for the provision of multiple ecosystem services and the resilience of the system to environmental changes, respectively. For this, we simulated the restoration of 24 riparian woody communities of the Brazilian Cerrado. Using the species pool of each original local community, we ran restoration simulations for gradually increasing species richness until reaching the total species richness of the original local community. We computed FRic and FR for each simulated restoration community using the traits specific leaf area, maximum plant height and seed mass. Our simulation results indicate that multiple ecosystem services could be restored with an average of 66% of the species of the original community. Moreover, an average of 59% of the species would be needed to restore communities resilient to environmental changes. Our approach contributes to solving one of the major challenges of ecological restoration, which is defining how many and which species should be used to achieve functional targets. We believe this approach can help in projects of restoration by enabling restoration practitioners to select minimum alternative sets of species that optimize the provision of multiple ecosystem services in a resilient restored ecosystem.

## Introduction

The delivery of ecosystem services, i.e. benefits provided by nature of relevant interest to human beings (MEA 2005), is one of the main objectives for restoring degraded areas (Rey-Benayas *et al.* 2009; Wainwright *et al.* 2018). For a restored ecosystem to deliver services, it is necessary to guarantee the recovery of its functioning in concert with the reassembly of native species. Ecosystem functioning can be assessed through functional structure (Laughlin 2014), which is well represented by the distribution of species and their abundances in a multivariate space defined by their functional traits (Villéger, Mason & Mouillot 2008). Functional traits are morpho-pheno-physiological characteristics of individuals that affect their fitness and performance in the environment (Violle *et al.* 2007). Functional traits can influence ecosystem functioning because they respond to environmental changes and disturbances as well as affect ecosystem processes (Lavorel & Garnier 2002; McGill *et al.* 2006). Ecosystem processes and functions are regulated by functional traits and promote ecosystem services, such as the supply of water, wood, pollination, weather control, ecotourism, nutrient cycling and soil formation (MEA 2005). Therefore, it is important to target ecosystem services mediated by functional traits in addition to species composition when restoring a degraded area (Montoya, Rogers & Memmott 2012; Laughlin 2014; Ostertag *et al.* 2015; Cadotte *et al.* 2015).

Functional trait-based approaches may be more appropriate than taxonomic approaches for restoration for recovering ecosystem services, especially in species-rich ecosystems (Carlucci et al. In press). Species composition is variable during community succession and has been shown to be less predictable as a restoration success measure than functional trait-based measures (Laughlin *et al.* 2017). Further, communities with different taxonomic composition may present similar functions (Laughlin *et al.* 2017). Meeting the major recommendation of using high species diversity in restoration has been a big challenge, especially in megadiverse ecosystems like tropical forests (Rodrigues *et al.* 2009; Brancalion *et al.* 2010). A major issue to follow this recommendation is the usual lack of native species as seeds or saplings in the market, which is especially challenging in megadiverse tropical countries (Stanturf, Palik & Dumroese 2014; Silva *et al.* 2017). Functional trait-based restoration is a recent approach that might help solve this lack of plant materials for restoration. This approach focus on the use of functional trait to target ecosystem services in restoration rather than aiming to recover species composition of a reference ecosystem per se (Laughlin 2014; Rayome *et al.* 2019). A functional trait-based approach for restoration enables the use of characteristics measured in the species that can be proxies for ecosystem targets (Montoya, Rogers & Memmott 2012; Laughlin 2014; Cadotte *et al.* 2015).

The diversity of functional traits positively affects community functioning via complementary resource use, which can mediate stable species coexistence (Loreau & Hector 2001; Morin *et al.* 2011; Lohbeck *et al.* 2015; Chen *et al.* 2016). Species usually exhibit ecological differences, either in the way they acquire resources or respond to abiotic conditions, enabling their coexistence (Turnbull *et al.* 2016). The trait-based approach enables a more mechanistic assessment of the relationship between the biota and ecosystem functioning than the taxonomical approach (Petchey & Gaston 2006). Multiple functional traits may provide diverse ecosystem functions and contribute to several ecosystem processes provisioning some ecosystem services (de Bello *et al.* 2010). Therefore, an index of functional richness, i.e. a measure of the amount of information an ecological community holds in terms of multiple traits carried by their species, may serve as an indicator of the degree in which multiple ecosystem services are provided by that community. Considering that functional trait diversity can be interpreted as diversity of ecosystem functions (Cadotte, Carscadden & Mirotchnick 2011), the functional richness (FRic) index (Cornwell, Schwilk & Ackerly 2006; Villéger, Mason & Mouillot 2008) serves this purpose. FRic measures the amount of trait information in the multivariate space occupied by co-occurring species.

The identification of long-run strategies of species persistence is important in restoration projects (SER 2004). According to the insurance hypothesis, more diverse communities have higher probability of persisting to environmental changes in the long run, because of possible increased diversity in response to environmental pressures (Yachi & Loreau 1999; Elmqvist *et al.* 2003). The relationship between functional diversity and stability is then accounted for by the FRic index. Nevertheless, the insurance hypothesis also predicts that among functionally diverse communities, a community with high functional redundancy should be more stable (Pillar *et al.* 2013). Highly redundant communities would be more stable because several species would provide the same function, thereby reducing the negative effect of species extinction on the loss of ecosystem functions (Fonseca & Ganade 2001). In this sense, the index of functional redundancy (FR; de Bello et al. 2007; Ricotta et al. 2016) may be used as a measure of resilience of the system against environmental changes and disturbances (Pillar *et al.* 2013; Ricotta *et al.* 2020).

Recently, Laughlin *et al.* (2018) proposed a framework to select species for restoration based on functional diversity. They used Rao functional diversity (Botta-Dukát 2005), which measures mean trait dispersion among co-occurring species as a proxy for niche complementarity mediating ecosystem functioning, i.e. multiple ecosystem services. Although that study represents an important advancement for functional restoration, it does not permit the selection of a minimum set of species for maximizing multiple ecosystem services and stability.

In the present study, our aim was to fill this gap by providing a trait-based framework that enables restoration practitioners to select a minimal set of species that maximize multiple ecosystem services and resilience to environmental changes and disturbances. For this, we simulated the restoration of communities, starting with a small subset of species from the original community and gradually adding species until the total species richness of the original community was reached. We used FRic index as a proxy for the provision of multiple services by the restored community, under the assumption that higher FRic corresponds to more services provided by the ecosystem. Moreover, we assessed FR at each simulated step of the restoration as a proxy of resilience of the system to changes in the environment.

We used as the study system riparian woody communities of the Brazilian Cerrado, which are species-rich tropical vegetation often threatened by agriculture, cattle ranching, and logging (Baptista-Maria *et al.* 2009). Riparian plant communities influence water bodies by modifying their microclimate through shading and by protecting the riverbanks against erosion, and the terrestrial ecosystem by affecting soil formation through organic matter input and soil protection, which makes the system rather complex and dynamic (Gregory *et al.* 1991; Naiman & Décamps 1997). We discuss how many species on average are needed to maximize multiple ecosystem services and resilience in the restoration of Cerrado riparian communities.

## Materials and Methods

### Compilation of data of riparian woody communities

We used Cerrado riparian communities composed by arboreal and arborescent species as a study system to test the trait-based framework proposed. The arboreal and arborescent species (hereafter, “woody species”, for simplification) included trees, treelets, tall shrubs and palms. We compiled data of riparian woody communities from scientific articles using Google Scholar and Web of Science databases (accessed from August to September 2016). For the search, we used the following keywords in English: (“riparian vegetation” OR “gallery forest”) AND (“phytosociology” OR “floristic survey”) AND “cerrado”. In Portuguese, we used: (“vegetação ripária” OR “mata de galeria” OR “mata ciliar”) AND (“fitossociologia” OR “inventário florístico”) AND “cerrado”. We compiled one or more lists of woody species reported in each study and recorded the respective location and geographical coordinates. We obtained a total of 68 studies containing information of 101 local communities and 1,953 woody species.

We validated and standardized botanical nomenclature according to the Brazilian Flora 2020 (http://floradobrasil.jbrj.gov.br/; accessed on November 2017), using the function “get.taxa”, “flora” package in R (R Core Team 2017). The nomenclature of species not resolved by the Brazilian Flora were validated by consulting the The Plant List database (http://www.theplantlist.org/; accessed on November 2017).

### Functional traits

We used functional traits that characterize major dimensions of plant ecological strategies in distinct environments (Westoby *et al.* 2002). For this, we used the LHS scheme (leaf-height-seed) proposed by Westoby (1998). In LHS, the traits specific leaf area (SLA), maximum height (H_max_) and seed mass (SM) characterize several important aspects of plant strategies, like competition, responses to disturbances and environmental stress, and tradeoffs in resource use and plant allometry (Westoby *et al.* 2002).

SLA is a key trait in the leaf economics spectrum (Wright *et al.* 2004; Reich 2014). High SLA values are related to fast leaf turnover, high rate of photosynthesis and fast growth and advantages for light competition, while low SLA usually results in higher leaf lifespan, lower rate of photosynthesis, and a slower growth, but higher environmental stress resistance (Westoby 1998). H_max_ is a good indicator of competitive ranks, because taller plants have advantage in light competition in humid ecosystems that can support high leaf area (Westoby *et al.* 2002; Laughlin 2014). SM is a good indicator of tradeoffs between resource allocation to seed production in mature plants and seedling germination and establishment, and may represent contrasts between growth and survival or colonization and competition strategies (Turnbull, Rees & Crawley 1999; Poorter *et al.* 2008). SM is usually negatively correlated with seed dispersal ability (Tamme *et al.* 2014). Species with large seeds have higher energy reserves, so that their saplings are able to persist in poorer soils (Westoby 1998). On the other hand, species producing smaller seeds are usually able to produce large numbers of seeds, which increase the probability of dispersing to and colonizing a higher number of places (Westoby 1998). LHS traits ultimately mediate ecosystem processes and functions important to ecosystem services provided by riparian woody communities (Table 1).

**Table 1.**
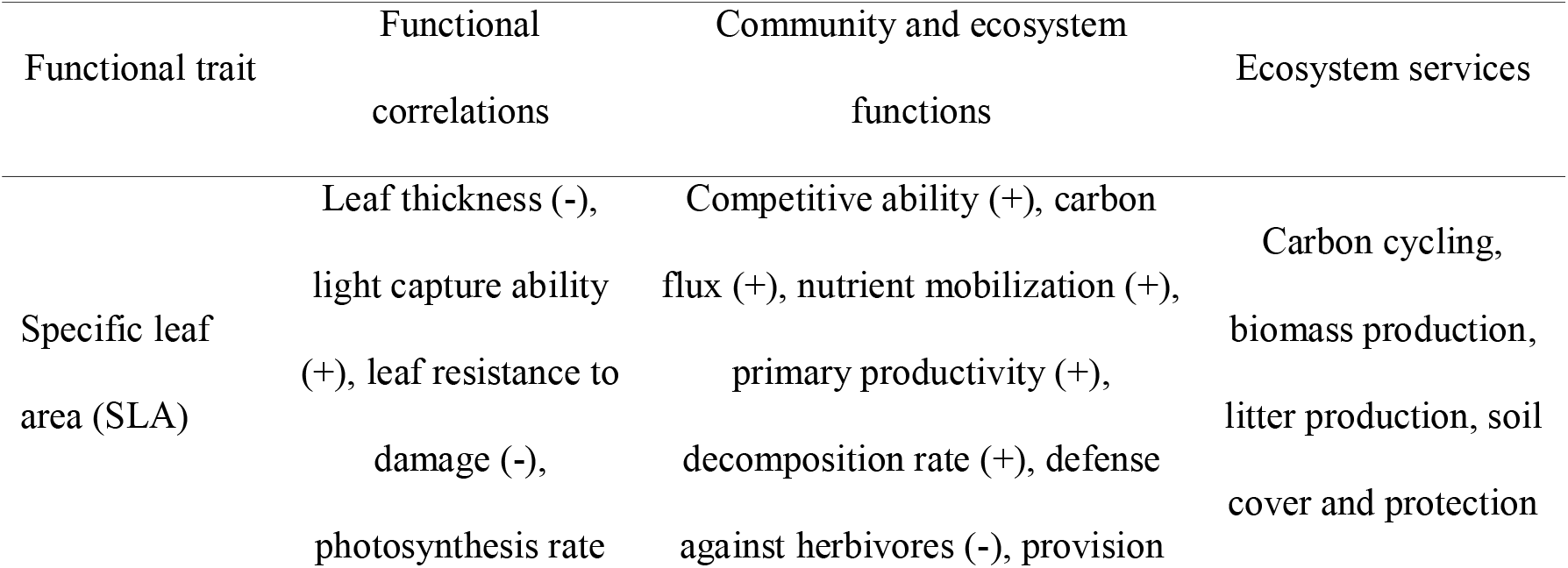
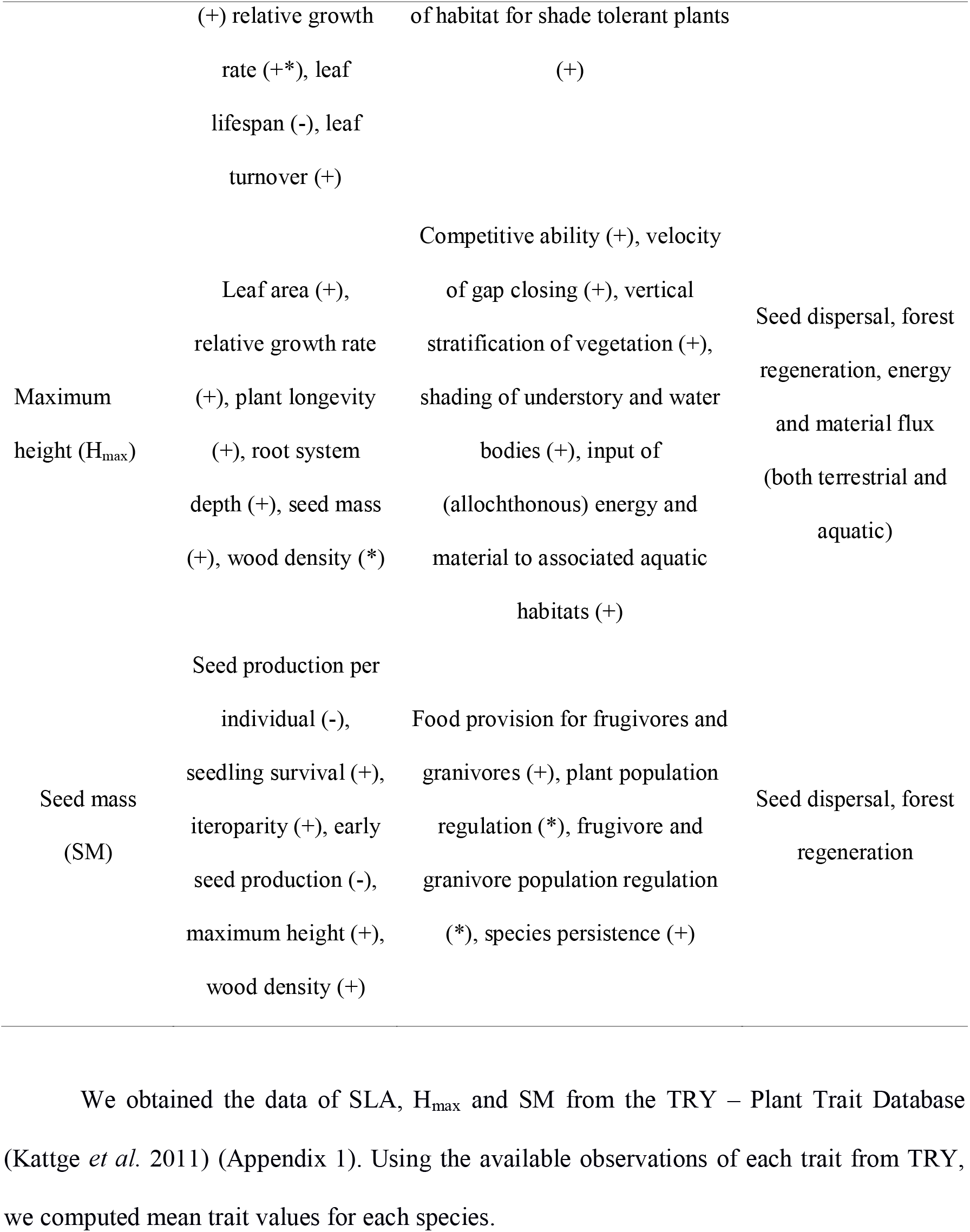
Functional traits, their correlations with other traits and their relationship with ecosystem functions and services. Positive (+) and negative (−) relationships are indicated. Complex relationships are indicated with an asterisk (*). The relationships indicated here are based on the following studies: Westoby (1998), de Bello *et al.* (2010); Moles (2018).

We obtained the data of SLA, H_max_ and SM from the TRY – Plant Trait Database (Kattge *et al.* 2011) (Appendix 1). Using the available observations of each trait from TRY, we computed mean trait values for each species.

### Selection of communities based on available trait information

We selected riparian communities that had at least 40% of species with TRY data available for the three traits (SLA, H_max_ and SM), which resulted in a total of 33 communities and a total of 559 species. We used two approaches to increase the number of species with trait information available. First, we performed an additional search to obtain missing species trait information in the specialized literature (Appendix 2). We used trait mean values whenever a species trait information was available in more than one source, which still resulted in a total of 139 species with missing traits information (24,87%). Second, we carried out a data imputation for species still missing information by using trait mean at the genus level, or by using trait mean at family level for genera with information for only one species. After the imputation, there were 39 species without trait information (7,2%). We selected communities with at least 90% of species with information for SLA, H_max_ and SM, and excluded species lacking trait information from the analysis. This whole procedure resulted in 24 communities (Appendix 3), with seven to 130 species, and 318 species in total (Appendix 4).

### Trait-based descriptors of functional structure

We evaluated two aspects of functional structure of the community: functional richness (FRic), as a proxy for multiple services provided by the restored ecosystem, and functional redundancy (FR), as a proxy of resilience to disturbances and environmental changes. Functional richness can be measured as a trait hypervolume (Cornwell, Schwilk & Ackerly 2006; Villéger, Mason & Mouillot 2008), in which a minimum convex volume connects externally species distributed in a trait multivariate space. We used the functional redundancy index described by Ricotta *et al.* (2016), originally proposed by de de Bello *et al.* (2007). FR is the complement of mean functional diversity, calculated by subtracting the Rao index value (Q; Botta-Dukát 2005) from the species diversity value computed using the Simpson index (D), that is, FR = D - Q. Ricotta *et al.* (2016) suggests to divide FR by the Simpson index, to standardize the index: FR = (D - Q)/D. This standardization enables the comparison of FR values between communities with varying species richness. We standardized trait data to zero mean and unit variance (within trait variables) and used Euclidean distances between species to compute both FRic and FR. We used only species occurrences (not abundances) to compute D and Q, to make FR comparable with FRic, which by definition does not use abundances.

### Simulation of community restoration while maximizing ecosystem services and resilience

To find the minimal sets of species for restoration that maximize the delivery of multiple services and a resilient ecosystem, we developed an algorithm to simulate local community reassembly, which looks for minimal sets of species with the most functional richness and redundancy. The algorithm works by generating species subsets of the original local community varying in species richness. The algorithm takes as input the species found in the original local community, simulating its restoration after the loss of the original community (Fig 1). In the first step, random communities are generated, with increasing species richness, from three to the total number of species of the original local community. For each species richness, 100 communities are generated by random selection of species from the total original pool of that local community. In other words, an original local community is subsetted to three random species a hundred times, then is subsetted to four random species a hundred times and so on until total species richness of the original local community is reached. FRic and FR are computed for each restored community.

**Figure 1.**
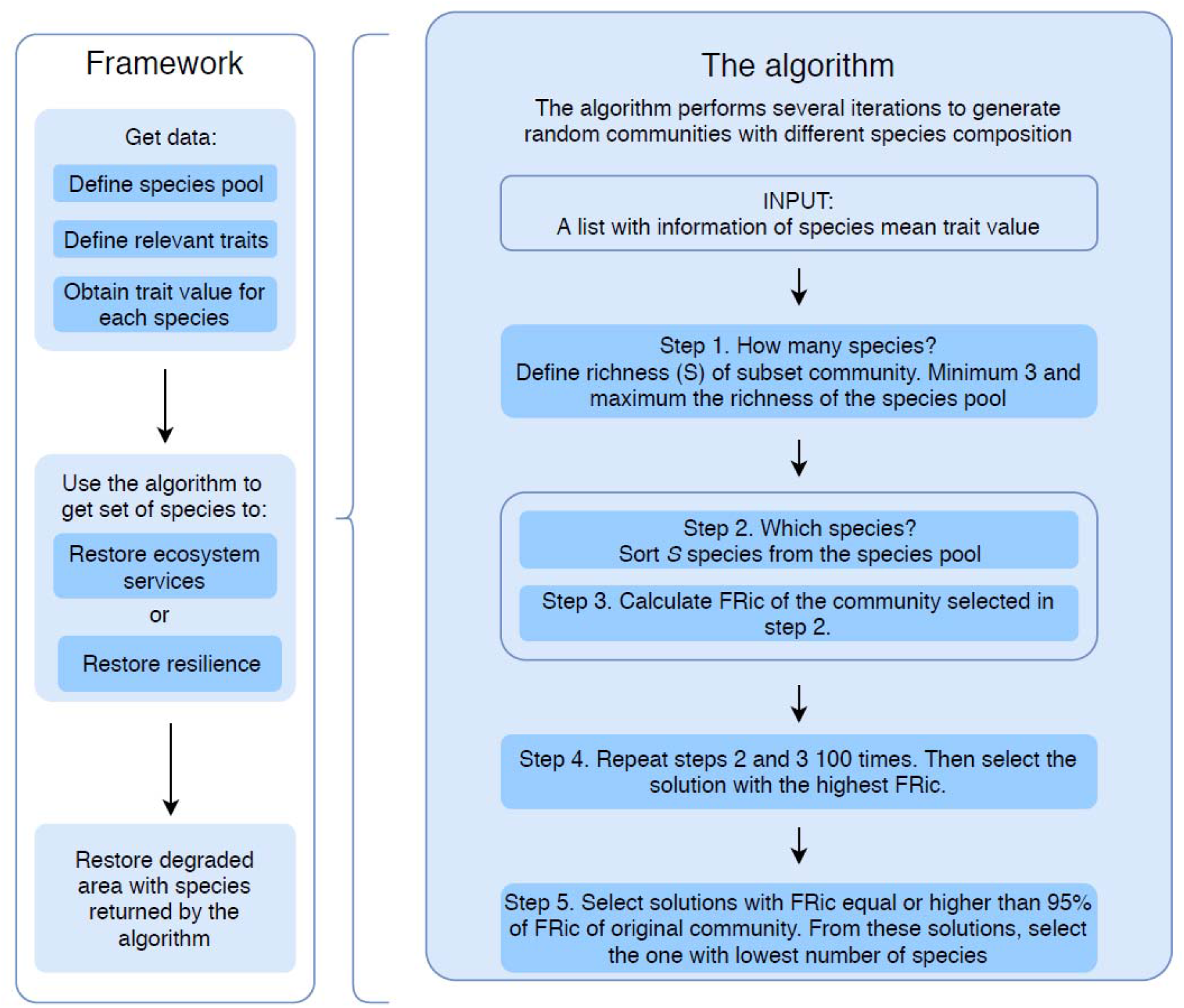
Steps of the framework to select species to maximize the functional richness (FRic) and functional redundancy (FR). First, the species pool and the relevant traits must be defined. In this study, we used the set of species of the original local community as a reference for restoration. Then, information of trait value for each species must be accessed. The algorithm uses this information to get a set of species that can maximize provision of ecosystem services or ecosystem resilience, with minimum species richness (S). The algorithm takes as input a vector with the information of species trait values. In the first step, it generates communities with different S values, starting from three species to the number of species in the original species pool. For each S value, S random species are selected, and FRic and FR values are computed. This process is repeated 100 times for each S. Finally, communities with the highest FRic and FR are selected for each S. Then, the communities with FRic higher than 95% of the original community are kept. From these solutions, the community with the lowest number of species is selected. FRic, FR and species composition of this community are returned by the algorithm.

For each species richness, the community with higher FRic is selected (FRicmax), as well as the community with higher FR (FR_max_). Then, solutions with FRic_max_ and FR_max_ equal to or higher than 95% of the FRic of original community are selected. Among these, the solution with the lowest number of species is selected, resulting in two solutions, one that maximizes FRic and the other that maximizes FR. We considered that these solutions maximize FRic and FR because they recover 95% of the values of these measures for the original community, with the lowest species richness possible.

### Data analyses

The number of species and the set of species used in the algorithm to maximize FRic can differ from those used to maximize FR. The ideal scenario would be that the same number and set of species could maximize both FRic and FR. We evaluated if, for the riparian communities selected, there was a significant difference between the number of species to maximize FRic and the number of species to maximize FR, using a paired *t*-test.

Another difference that may arise is that, while maximizing FRic, communities could have lower FR values than those when FR was maximized and vice-versa. First, we evaluated if that was the case by comparing FR of the community with the highest FRic was significantly different from the FR of the community when FR was maximized. For this, we compared FR of the community with highest FRic with the FR_max_, that is, the FR of the community in the FR threshold, using a paired *t*-test. Second, we evaluated if the FRic of the community with the highest FR was significantly different from the maximum FRic of the communities that maximized this parameter (FRicmax). For testing this, we performed Wilcoxon test, because data was not normally distributed. All other data was normally distributed and satisfied *t*-test assumptions.

All analyses were carried using the statistical software R (R Core Team 2017).

## RESULTS

The proportion of species to maximize FRic was higher than the proportion to maximize FR (Fig. 2A). Our simulations (Appendix 5) indicated that the presence, on average, of 66% of the original species is necessary to recover 95% of FRic of the original riparian woody communities, enabling the provision of multiple ecosystem services. On the other hand, our simulations (Appendix 6) indicated that to recover 95% of FR of the original riparian woody communities, and thereby guarantee community resilience, on average 59% of the original species are required.

**Figure 2.**
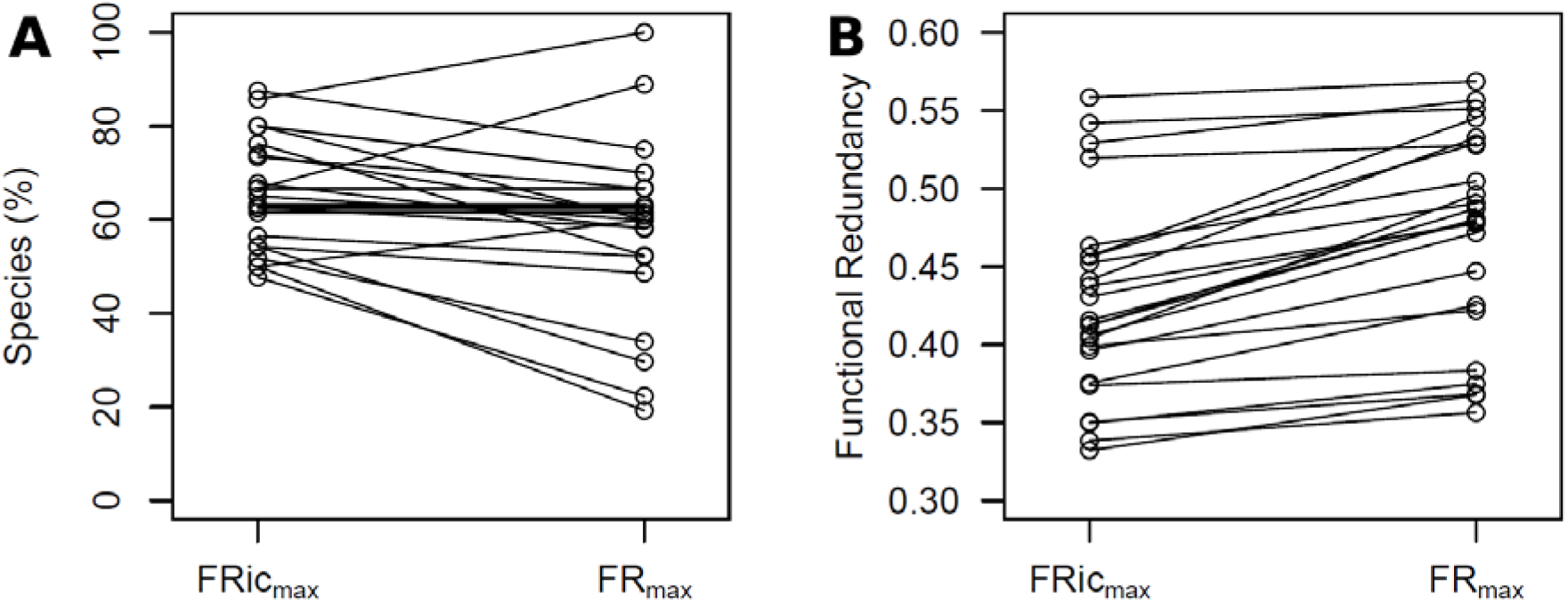
(A) Species proportion of the selected solution to maximize FRic and FR of the 24 riparian woody communities. Lines connect the solutions of each community. Paired *t-*test: *t* = 2.84, *d.f.* = 23, *p* = 0.009. (B) FR of the selected solution to maximize FRic and FR of the 24 riparian woody communities. Lines connect the solutions of each community. Paired *t-*test: *t* = −7.98, *d.f.* = 23,*p* < 0.001.

The FR of communities that maximized FRic was, on average, 4% lower than in the communities that maximized FR (Fig. 2B). The FRic of species that maximizes FR was, on average, 43% less than FRic of the community that maximizes FRic *(W* = 554, *p* < 0.001, Appendix 7).

The resulting curves of FRic and FR vs. increasing species richness in simulated restoration of each riparian woody community are available in Appendices 6 and 7.

## DISCUSSION

### A few or lots of: how many species should we use in restoration?

Restoration ecology has traditionally focused on recovering species composition of an originally lost community, or at least of a reference community harboring high similarity with the original community (SER 2004). The traditional restoration has been widely used since the foundations of restoration ecology (Palmer, Zedler & Falk 2016), especially in temperate ecosystems in developed countries (Kollmann *et al.* 2016; Wilson *et al.* 2016). However, in tropical, usually developing countries, it became hard to accomplish the traditional restoration, because of the high species richness, lack of knowledge on the original species composition of lost communities, biodiversity knowledge shortfalls, and lack of species in the market (Karlsson, Srebotnjak & Gonzales 2007; Stanturf, Palik & Dumroese 2014; Hortal *et al.* 2015; Silva *et al.* 2017). Despite these several issues, there is isolated evidence that after planting high number of native species (>50 species) in megadiverse tropical ecosystems, restoration programs succeeded not only in recovering species composition of reference areas, but also functional diversity and ecosystem services (Rodrigues *et al.* 2009). Nevertheless, accomplishing high species richness restoration is a major challenge to most tropical regions, because it is expensive and demands high scientific development.

To avoid a dichotomical discussion on whether we need to plant few or many species or a fuzzy discussion on how many and which species species are necessary to recover ecosystem services, we need a clear theory-based framework. Restoration ecology has increasingly incorporated advances of ecological theory into restoration practices (Wainwright *et al.* 2018). More specifically, trait-based frameworks for restoring ecosystem services have advanced in important ways towards a more predictive restoration (Laughlin 2014; Ostertag *et al.* 2015; Laughlin *et al.* 2018; Rayome *et al.* 2019). Despite the advancements, these frameworks do not inform how many species are necessary to achieve a maximum of ecosystem service provision and resilience with a minimal set of species. Our framework was built exactly to fill this gap and should help optimizing restoration, especially in megadiverse ecosystems.

### Changing the focus from species recomposition to functional restoration

The importance of considering the recovery of ecosystem services with ecological restoration of ecosystem is not new, but only more recently clear quantitative frameworks to restore services using functional traits started to be created (e.g. Laughlin 2014; Laughlin *et al.* 2018). While these frameworks represent important advancements and may be used to monitor recovery of functional structure (e.g. Rosenfield *et al.* 2017) and even be used to project restoration since its beginning, our framework represents further advancements in important ways.

First, we propose a way to maximize multiple ecosystem services in restoration using a minimal set of species. Laughlin *et al.* (2018) uses Rao index as a measure of functional diversity, focusing on the niche complementarity theory. Given that Rao diversity is the average of pairwise trait differences between cooccurring species (Botta-Dukat 2005), Laughlin *et al.* (2018) approach could not be used to maximize multiple ecosystem services. While Rao diversity index averages species differences, thereby causing loss of information on the trait space, functional richness (FRic) index has the property of quantifying the amount of trait information defined by coccurring species (Villéger, Mason & Mouillot 2008). Once a set of functional traits related to multiple ecosystem services are chosen, FRic might be interpreted against species richness as a measure of amount of ecosystem services provided by different species sets. Our framework thus enable to predict how many and which species will be necessary to reach a maximum of ecosystem services provided by the community to be restored.

Second, our framework also accounts for the resilience of the restored community, which no other exisiting functional trait-based framework for restoration has attempted to do before, to our knowledge. Considering that we live in a rapidly changing world, only maximizing ecosystem services is not enough to pursue a stable restored community. The insurance hypothesis (Yachi & Loreau 2000) considers that having functionally similar species coocurring would make the community more resilient to disturbances and environmental changes, because the loss of a certain species would be balanced by a functionally similar species remaining in the system. The use of a functional redundancy (FR) index enabled that we assessed redundancy in tandem with FRic, i.e. using the same methods for standardization of trait data and computation of trait distances between species.

### Should we maximize ecosystem services or stability or both?

Our results indicated that the set of species that maximized FRic in general did not present enough FR. In other words, it takes more species to recover multiple ecosystem services than resilience in the restored system. Therefore, restoration practitioners should also consider the set of species that maximize FR to assure there will be sufficient similar species in the system to face potential future disturbances and environmental changes. Ideally the practitioner should add those species that maximize FR but that were not present in the solution that maximized FRic. By doing so, the final community will have all the species that are needed to maximize both parameters. Using only the set of species that maximixed FR is not recommended, because it presented, on average, 43% less FRic than it could reach. Therefore, after running the algorithm, we should try and maximize both multiple services and stability of the system to be restored.

### Defining reference species pools based on areas or regions

In this study, we used the set of species presented in each original local community as the reference for restoration simulations. In the real word, information on the species composition of original communities is very rare and might not be the best option at hand considering the history of anthropogenic impacts ecosystems have been subject to (Thorpe & Stanley 2011). One approach to solve this problem is to use a species pool of the dark diversity, i.e. the species from the regional pool that are absent in some locations but that have potential to occur under certain environmental conditions (Pärtel, Szava-Kovats & Zobel 2011). This set of species could be used as input in the algorithm to verify the functional structure that reaches higher FRic and FR with the lowest number of species.

### Advancing biodiversity knowledge to accomplish broad-scale restoration

Trait information is essential to the application of our framework, but it can be hardly available for some organisms and ecosystems (Cornwell *et al.* 2019). Traits can be difficult to measure in regions with difficult access, few scientists, scarce resources, many rare species (Hortal *et al.* 2015). The system used in our study demonstrates the challenge of obtaining trait information to apply trait-based frameworks. We obtained information for less than 45% of species (from the initial community data compilation) using a worldwide comprehensive plant trait database. It is important to prioritize and define strategies to guide efforts to collect trait data, integrating technical and scientific knowledge with regional demands for restoration.

Lack of species as seeds in the market or as seedlings in nurseries is common particularly in megadiverse, developing countries (Stanturf, Palik & Dumroese 2014), and may also represent a challenge to apply our framework in some regions. In Brazil, for instance, higher availability of native species is found in the most economically-developed regions of the country, while restoration practitioners of other regions have difficulty in finding seeds or saplings for most of their native species (Silva *et al.* 2017). The species list generated by our algorithm could be used to identify species that are lacking in the market, and then support prioritization of biodiversity studies towards poorly known species and regions (Schmidt *et al.* 2019).

### How to deal with species abundances once a species set was defined?

Our framework was built to find the number of species and which species to use in restoration, which is a first step in restoration projects. Once the species list is defined, the restoration practitioner needs to know how many individuals of each species to plant, considering that the species abundances are also important to the provision of ecosystem services (Grime 1998; Winfree *et al.* 2015). For some regions, it is possible that this information is available in regional restoration manuals (e.g. Durigan *et al.* 2011), which then may be used to adjust the abundance of species returned by our algorithm. However, when species abundance information is missing, which is common, we suggest the use of the framework by Laughlin (2014), which enables to define species abundances according to ecosystem service targets. Therefore, the species list provided by the algorithm can be used as a starting point in restoration programs to reduce the number of species required to maximize ecosystem services and resilience; with the set of species at hand, the practitioner then may define species abundances, either by using specific information from regional restoration protocols or other specific trait-based frameworks.

## CONCLUSIONS

Computer simulations can give predictive solutions to environmental problems using ecological theories. Our framework provides minimum sets of species to maximize the delivery of multiple ecosystem services in a resilient restored ecosystem. With our framework, we hope to contribute to advance application of ecological theory into ecological restoration.

## AUTHORS’ CONTRIBUTIONS

P.K.T. and M.B.C. conceived the ideas; P.K.T. compiled the data; P.K.T., A.G.C., A.M.M., and M.B.C. analyzed the data and led the writing; N.J.B.K., A.G.M., P.H., S.C.M., Ü.N., V.D.P., and E.S. contributed with trait data via TRY; all authors contributed to revise the drafts and gave final approval for publication.

## ACKNOWLEDGEMENTS

This study was financed by Coordenação de Aperfeiçoamento de Pessoal de Nível Superior - Brazil (CAPES) - Finance Code 001 – via a postdoctoral fellowship to M.B.C., a MSc fellowship to P.K.T., and PhD fellowships to A.G.C. and A.M.M.. This paper is a contribution of the INCT in Ecology, Evolution and Biodiversity Conservation founded by MCTIC/CNPq (grant #465610/2014-5) and FAPEG (grant #201810267000023). We thank Marcus Vinicius Cianciaruso for useful discussions during the course of this study.

## DATA ACCESSIBILITY

This study used data provided by the TRY database (https://www.try-db.org) - a network of vegetation scientists headed by Future Earth, the Max Planck Institute for Biogeochemistry, and iDiv providing open access to plant trait data.

## REFERENCES

de Bello, F., Lavorel, S., Díaz, S., Harrington, R., Cornelissen, J.H.C., Bardgett, R.D., … Harrison, P.A. (2010) Towards an assessment of multiple ecosystem processes and services via functional traits. Biodiversity and Conservation, 19, 2873–2893.

de Bello, F., Lepš, J., Lavorel, S. & Moretti, M. (2007) Importance of species abundance for assessment of trait composition: an example based on pollinator communities. Community Ecology, 8, 163–170.

Botta-Dukát, Z. (2005) Rao’s quadratic entropy as a measure of functional diversity based on multiple traits. Journal of vegetation science, 16, 533–540.

Brancalion, P.H.S., Rodrigues, R.R., Gandolfi, S., Kageyama, P.Y., Nave, A.G., Gandara, F.B., … Tabarelli, M. (2010) Instrumentos legais podem contribuir para a restauração de florestas tropicais biodiversas. Revista Árvore, 34, 455–470.

Cadotte, M.W., Arnillas, C.A., Livingstone, S.W. & Yasui, S.-L.E. (2015) Predicting communities from functional traits. Trends in Ecology & Evolution, 30, 510–511.

Cadotte, M.W., Carscadden, K., & Mirotchnick, N. (2011). Beyond species: functional diversity and the maintenance of ecological processes and services. Journal of Applied Ecology, 48, 1079–1087.

Carlucci, M. B., Brancalion, P. H., Rodrigues, R. R., Loyola, R., & Cianciaruso, M. V. (In press). Functional traits and ecosystem services in ecological restoration. Restoration Ecology, https://doi.org/10.1111/rec.13279.

Chen, Y., Wright, S.J., Muller-Landau, H., Hubbell, S., Wang, Y. & Yu, S. (2016) Positive effects of neighborhood complementarity on tree growth in a Neotropical forest. Ecology, 97, 776–785.

Cornwell, W.K., Pearse, W.D., Dalrymple, R.L. & Zanne, A.E. (2019) What we (don’t) know about global plant diversity. Ecography, 42, 1819–1831.

Cornwell, W.K., Schwilk, D.W. & Ackerly, D.D. (2006) A trait-based test for habitat filtering: convex hull volume. Ecology, 87, 1465–1471.

Durigan, G., Melo, A.C.G., Max, J.C.M., Vilas Boas, O., Contieri, W.A. & Ramos, V.S. (2011) Manual para recuperação da vegetação de cerrado, 3rd ed. SMA, São Paulo.

Elmqvist, T., Folke, C., Nyström, M., Peterson, G., Bengtsson, J., Walker, B. & Norberg, J. (2003) Response diversity, ecosystem change, and resilience. Frontiers in Ecology and the Environment, 1, 488–494.

Fonseca, C.R. & Ganade, G. (2001) Species functional redundancy, random extinctions and the stability of ecosystems. Journal of Ecology, 89, 118–125.

Gregory, S.V., Swanson, F.J., McKee, W.A. & Cummins, K.W. (1991) An ecosystem perspective of riparian zones. BioScience, 41, 540–551.

Grime, J.P. (1998) Benefits of plant diversity to ecosystems: immediate, filter and founder effects. Journal of Ecology, 86, 902–910.

Hortal, J., de Bello, F., Diniz-Filho, J.A.F., Lewinsohn, T.M., Lobo, J.M. & Ladle, R.J. (2015) Seven shortfalls that beset large-scale knowledge of biodiversity. Annual Review of Ecology, Evolution, and Systematics, 46, 523–549.

Karlsson, S., Srebotnjak, T. & Gonzales, P. (2007) Understanding the North–South knowledge divide and its implications for policy: a quantitative analysis of the generation of scientific knowledge in the environmental sciences. Environmental Science & Policy, 10, 668–684.

Kattge, J., Díaz, S., Lavorel, S., Prentice, I.C., Leadley, P., Bönisch, G., … Wirth, C. (2011) TRY - a global database of plant traits. Global Change Biology, 17, 2905–2935.

Kollmann, J., Meyer, S.T., Bateman, R., Conradi, T., Gossner, M.M., de Souza Mendonça, M., … Weisser, W.W. (2016) Integrating ecosystem functions into restoration ecology-recent advances and future directions: Ecosystem functions in restoration ecology. Restoration Ecology, 24, 722–730.

Laughlin, D.C. (2014) Applying trait-based models to achieve functional targets for theory-driven ecological restoration. Ecology Letters, 17, 771–784.

Laughlin, D.C., Chalmandrier, L., Joshi, C., Renton, M., Dwyer, J.M. & Funk, J.L. (2018) Generating species assemblages for restoration and experimentation: a new method that can simultaneously converge on average trait values and maximize functional diversity. Methods in Ecology and Evolution, 9, 1764–1771.

Laughlin, D.C., Strahan, R.T., Moore, M.M., Fulé, P.Z., Huffman, D.W. & Covington, W.W. (2017) The hierarchy of predictability in ecological restoration: are vegetation structure and functional diversity more predictable than community composition? Journal of Applied Ecology, 54, 1058–1069.

Lavorel, S. & Garnier, É. (2002) Predicting changes in community composition and ecosystem functioning from plant traits: revisiting the Holy Grail. Functional ecology, 16, 545–556.

Lohbeck, M., Poorter, L., Martínez-Ramos, M. & Bongers, F. (2015) Biomass is the main driver of changes in ecosystem process rates during tropical forest succession. Ecology, 96, 1242–1252.

Loreau, M. & Hector, A. (2001) Partitioning selection and complementarity in biodiversity experiments. Nature, 412, 72–76.

McGill, B.J., Enquist, B.J., Weiher, E. & Westoby, M. (2006) Rebuilding community ecology from functional traits. Trends in Ecology & Evolution, 21, 178–185.

MEA - Millennium Ecosystem Assessment (2005) Ecosystems and Human Well-Being: Synthesis. Island Press, Washington, DC.

Moles, A.T. (2018) Being John Harper: Using evolutionary ideas to improve understanding of global patterns in plant traits. Journal of Ecology, 106, 1–18.

Montoya, D., Rogers, L. & Memmott, J. (2012) Emerging perspectives in the restoration of biodiversity-based ecosystem services. Trends in Ecology & Evolution, 27, 666–672.

Morin, X., Fahse, L., Scherer□Lorenzen, M. & Bugmann, H. (2011) Tree species richness promotes productivity in temperate forests through strong complementarity between species. Ecology Letters, 14, 1211–1219.

Naiman, R.J. & Décamps, H. (1997) The ecology of interfaces: Riparian zones. Annual Review of Ecology and Systematics, 28, 621–658.

Ostertag, R., Warman, L., Cordell, S. & Vitousek, P.M. (2015) Using plant functional traits to restore Hawaiian rainforest. Journal of Applied Ecology, 52, 805–809.

Palmer, M.A., Zedler, J.B. & Falk, D.A. (2016) Ecological theory and restoration ecology. Foundations of Restoration Ecology, pp. 3–26. Island Press, Washington, DC.

Pärtel, M., Szava-Kovats, R. & Zobel, M. (2011) Dark diversity: shedding light on absent species. Trends in Ecology & Evolution, 26, 124–128.

Petchey, O.L. & Gaston, K.J. (2006) Functional diversity: back to basics and looking forward. Ecology Letters, 9, 741–758.

Pillar, V.D., Blanco, C.C., Müller, S.C., Sosinski, E.E., Joner, F. & Duarte, L.D.S. (2013) Functional redundancy and stability in plant communities. Journal of Vegetation Science, 24, 963–974.

Poorter, L., Wright, S.J., Paz, H., Ackerly, D.D., Condit, R., Ibarra-Manríquez, G., … Wright, I.J. (2008) Are functional traits good predictors of demographic rates? Evidence from five Neotropical forests. Ecology, 89, 1908–1920.

R Core Team (2019) R: A Language and Environment for Statistical Computing. R Foundation for Statistical Computing, Vienna, Austria. URL https://www.R-project.org/.

Rayome, D., DiManno, N., Ostertag, R., Cordell, S., Fung, B., Vizzone, A., … Tate, R. (2019) Restoring Ecosystem Services Tool (REST): A Computer Program for Selecting Species for Restoration Projects Using a Functional-Trait Approach. United States Department of Agriculture, US Forest Service, Hilo.

Reich, P.B. (2014) The world-wide ‘fast-slow’ plant economics spectrum: a traits manifesto. Journal of Ecology, 102, 275–301.

Rey-Benayas, J.M., Newton, A.C., Diaz, A. & Bullock, J.M. (2009) Enhancement of biodiversity and ecosystem services by ecological restoration: a meta-analysis. Science, 325, 1121–1124.

Ricotta, C., Bello, F. de, Moretti, M., Caccianiga, M., Cerabolini, B.E.L. & Pavoine, S. (2016) Measuring the functional redundancy of biological communities: a quantitative guide. Methods in Ecology and Evolution, 7, 1386–1395.

Rodrigues, R.R., Lima, R.A.F., Gandolfi, S. & Nave, A.G. (2009) On the restoration of high diversity forests: 30 years of experience in the Brazilian Atlantic Forest. Biological Conservation, 142, 1242–1251.

Rosenfield, M.F. & Müller, S.C. (2017) Predicting restored communities based on reference ecosystems using a trait-based approach. Forest Ecology and Management, 391, 176–183.

Schmidt, I.B., Urzedo, D.I. de, Piña□Rodrigues, F.C.M., Vieira, D.L.M., Rezende, G.M. de, Sampaio, A.B. & Junqueira, R.G.P. (2019) Community-based native seed production for restoration in Brazil – the role of science and policy. Plant Biology, 21, 389–397.

SER (2004) The SER International Primer on Ecological Restoration. Society for Ecological Restoration International Science & Policy Working Group.

Silva, A.P.M. da, Schweizer, D., Marques, H.R., Teixeira, A.M.C., Santos, T.V.M.N. dos, Sambuichi, R.H.R., … Brancalion, P.H.S. (2017) Can current native tree seedling production and infrastructure meet an increasing forest restoration demand in Brazil? Restoration Ecology, 25, 509–515.

Stanturf, J.A., Palik, B.J. & Dumroese, R.K. (2014) Contemporary forest restoration: A review emphasizing function. Forest Ecology and Management, 331, 292–323.

Tamme, R., Götzenberger, L., Zobel, M., Bullock, J.M., Hooftman, D.A.P., Kaasik, A. & Pärtel, M. (2014) Predicting species’ maximum dispersal distances from simple plant traits. Ecology, 95, 505–513.

Thorpe, A.S. & Stanley, A.G. (2011) Determining appropriate goals for restoration of imperilled communities and species: Defining appropriate restoration targets. Journal of Applied Ecology, 48, 275–279.

Turnbull, L.A., Isbell, F., Purves, D.W., Loreau, M. & Hector, A. (2016) Understanding the value of plant diversity for ecosystem functioning through niche theory. Proceedings of the Royal Society B: Biological Sciences, 283, 20160536.

Turnbull, L.A., Rees, M. & Crawley, M.J. (1999) Seed mass and the competition/colonization trade-off: a sowing experiment. Journal of Ecology, 87, 899–912.

Villéger, S., Mason, N.W. & Mouillot, D. (2008) New multidimensional functional diversity indices for a multifaceted framework in functional ecology. Ecology, 89, 2290–2301.

Violle, C., Navas, M.-L., Vile, D., Kazakou, E., Fortunel, C., Hummel, I. & Garnier, E. (2007) Let the concept of trait be functional! Oikos, 116, 882–892.

Wainwright, C.E., Staples, T.L., Charles, L.S., Flanagan, T.C., Lai, H.R., Loy, X., … Mayfield, M.M. (2018) Links between community ecology theory and ecological restoration are on the rise. Journal of Applied Ecology, 55, 570–581.

Westoby, M. (1998) A leaf-height-seed (LHS) plant ecology strategy scheme. Plant and Soil, 199, 213–227.

Westoby, M., Falster, D.S., Moles, A.T., Vesk, P.A. & Wright, I.J. (2002) Plant ecological strategies: Some leading dimensions of variation between species. Annual Review of Ecology and Systematics, 33, 125–159.

Wilson, K.A., Auerbach, N.A., Sam, K., Magini, A.G., Moss, A.St.L., Langhans, S.D., … Meijaard, E. (2016) Conservation research is not happening where it is most needed. PLoS Biology, 14, e1002413.

Winfree, R., Fox, J.W., Williams, N.M., Reilly, J.R. & Cariveau, D.P. (2015) Abundance of common species, not species richness, drives delivery of a real-world ecosystem service. Ecology Letters, 18, 626–635.

Wright, I.J., Reich, P.B., Westoby, M., Ackerly, D.D., Baruch, Z., Bongers, F., … Villar, R. (2004) The worldwide leaf economics spectrum. Nature, 428, 821–827.

Yachi, S. & Loreau, M. (1999) Biodiversity and ecosystem productivity in a fluctuating environment: The insurance hypothesis. Proceedings of the National Academy of Sciences of the United States of America, 96, 1463–1468.

