## Supplementary material for "How many and which species to plant? A multi-trait-based approach to select species to restore ecosystem services"

**APPENDIX 1.** References of the TRY data sources for species traits.

| **Data source** | **Publication** |
| --- | --- |
| ArtDeco Database | Cornwell, W. K., J. H. C. Cornelissen, K. Amatangelo, E. Dorrepaal, V. T. Eviner, O. Godoy, S. E. Hobbie, B. Hoorens, H. Kurokawa, N. Pérez-Harguindeguy, H. M. Quested, L. S. Santiago, D. A. Wardle, I. J. Wright, R. Aerts, S. D. Allison, P. van Bodegom, V. Brovkin, A. Chatain, T. V. Callaghan, S. Díaz, E. Garnier, D. E. Gurvich, E. Kazakou, J. A. Klein, J. Read, P. B. Reich, N. A. Soudzilovskaia, M. V. Vaieretti, and M. Westoby. 2008. Plant species traits are the predominant control on litter decomposition rates within biomes worldwide. Ecology Letters 11:1065-1071. |
| Cold Tolerance, Seed Size and Height of North American Forest Tree Species | unpub. |
| Costa Rica Rainforest Trees Database | unpub. |
| Costa Rican Tropical Dry Forest Trees | Jennifer S. Powers and Peter Tiffin 2012 Plant functional type classifications in tropical dry forests in Costa Rica: leaf habit versus taxonomic approaches. Functional Ecology 2010, 24, 927–936 doi: 10.1111/j.1365-2435.2010.01701.x |
| Crown Architecture Database | unpub. |
| Dispersal Traits Database | unpub. |
| ECOQUA South American Plant Traits Database | Muller, S. C., G. E. Overbeck, J. Pfadenhauer, and V. D. Pillar. 2007. Plant functional types of woody species related to fire disturbance in forest-grassland ecotones. Plant Ecology 189:1-14. |
| ECOQUA South American Plant Traits Database | Pillar, V. D. and E. E. Sosinski. 2003. An improved method for searching plant functional types by numerical analysis. Journal of Vegetation Science 14:323-332. |
| FAPESP Brazil Rainforest Database | Pillar, V. D. and E. E. Sosinski. 2003. An improved method for searching plant functional types by numerical analysis. Journal of Vegetation Science 14:323-332. |
| FAPESP Brazil Rainforest Database | unpub. |
| Fonseca/Wright New South Wales Database | Fonseca, C. R., J. M. Overton, B. Collins, and M. Westoby. 2000. Shifts in trait-combinations along rainfall and phosphorus gradients. Journal of Ecology 88:964-977. |
| Functional Traits for Restoration Ecology in the Colombian Amazon | unpub. |
| Global A, N, P, SLA Database | Reich, P. B., J. Oleksyn, and I. J. Wright. 2009. Leaf phosphorus influences the photosynthesis-nitrogen relation: a cross-biome analysis of 314 species. Oecologia 160:207-212. |
| Global Leaf Robustness and Physiology Database | Niinemets, U. 2001. Global-scale climatic controls of leaf dry mass per area, density, and thickness in trees and shrubs. Ecology 82:453-469. |
| Global Respiration Database | Reich, P. B., M. G. Tjoelker, K. S. Pregitzer, I. J. Wright, J. Oleksyn, and J. L. Machado. 2008. Scaling of respiration to nitrogen in leaves, stems and roots of higher land plants. Ecology Letters 11:793-801. |
| GLOPNET - Global Plant Trait Network Database | Wright, I. J., P. B. Reich, M. Westoby, D. D. Ackerly, Z. Baruch, F. Bongers, J. Cavender-Bares, T. Chapin, J. H. C. Cornelissen, M. Diemer, J. Flexas, E. Garnier, P. K. Groom, J. Gulias, K. Hikosaka, B. B. Lamont, T. Lee, W. Lee, C. Lusk, J. J. Midgley, M. L. Navas, U. Niinemets, J. Oleksyn, N. Osada, H. Poorter, P. Poot, L. Prior, V. I. Pyankov, C. Roumet, S. C. Thomas, M. G. Tjoelker, E. J. Veneklaas, and R. Villar. 2004. The worldwide leaf economics spectrum. Nature 428:821-827. |
| Hawaiian Leaf Traits Database | Peñuelas, J., J. Sardans, J. Llusia, S. Owen, J. Carnicer, T. W. Giambelluca, E. L. Rezende, M. Waite, and Ü. Niinemets. 2010. Faster returns on "leaf economics" and different biogeochemical niche in invasive compared with native plant species. Global Change Biology 16:2171-2185. |
| KEW African Plant Traits Database | Kirkup, D., P. Malcolm, G. Christian, and A. Paton. 2005. Towards a digital African Flora. Taxon 54:457-466. |
| LABDENDRO Brazilian Subtropical Forest Traits Database | unpub. |
| LBA ECO Tapajos: Leaf Characteristics and Photosynthesis | Tomas F. Domingues, Luiz A. Martinelli, James R. Ehleringer (2007) Ecophysiological traits of plant functional groups in forest and pasture ecosystems from eastern Amazonia, Brazil. Plant Ecol (2007) 193:101–112 DOI 10.1007/s11258-006-9251-z |
| Leaf and Whole Plant Traits Database | Shipley B., 2002. Trade-offs between net assimilation rate and specific leaf area in determining relative growth rate: relationship with daily irradiance, Functional Ecology(16) 682-689 |
| Leaf Biomechanics Database | Onoda, Y., M. Westoby, P. B. Adler, A. M. F. Choong, F. J. Clissold, J. H. C. Cornelissen, S. Diaz, N. J. Dominy, A. Elgart, L. Enrico, P. V. A. Fine, J. J. Howard, A. Jalili, K. Kitajima, H. Kurokawa, C. McArthur, P. W. Lucas, L. Markesteijn, N. Perez-Harguindeguy, L. Poorter, L. Richards, L. S. Santiago, Jr. E. Sosinski, S. Van Bael, D. I. Warton, I. J. Wright, S. J. Wright, and N. Yamashita. 2011 . Global patterns of leaf mechanical properties. Ecology Letters 14:301-312. |
| Leaf Physiology Database | Kattge, J., W. Knorr, T. Raddatz, and C. Wirth. 2009. Quantifying photosynthetic capacity and its relationship to leaf nitrogen content for global-scale terrestrial biosphere models. Global Change Biology 15:976-991. |
| Leaf Structure, Venation and Economic Spectrum | Blonder, B., Buzzard, B., Sloat, L., Simova, I., Lipson, R., Boyle, B., Enquist, B. (2012) The shrinkage effect biases estimates of paleoclimate. American Journal of Botany. 99.11 1756-1763 |
| Leaf Structure, Venation and Economic Spectrum | Blonder, B., Violle, C., Patrick, L., Enquist, B. Leaf venation networks and the origin of the leaf economics spectrum. Ecology Letters, 2011 |
| Neotropic Plant Traits Database | Wright, I. J., D. D. Ackerly, F. Bongers, K. E. Harms, G. Ibarra-Manriquez, M. Martinez-Ramos, S. J. Mazer, H. C. Muller-Landau, H. Paz, N. C. A. Pitman, L. Poorter, M. R. Silman, C. F. Vriesendorp, C. O. Webb, M. Westoby, and S. J. Wright. 2007. Relationships among ecologically important dimensions of plant trait variation in seven Neotropical forests. Annals of Botany 99:1003-1015. |
| Nutrient Resorption Efficiency Database | Vergutz, L., S. Manzoni, A. Porporato, R.F. Novais, and R.B. Jackson. 2012. A Global Database of Carbon and Nutrient Concentrations of Green and Senesced Leaves. Data set. Available on-line [http://daac.ornl.gov] from Oak Ridge National Laboratory Distributed Active Archive Center, Oak Ridge, Tennessee, U.S.A. http://dx.doi.org/10.3334/ORNLDAAC/1106 |
| Panama Leaf Traits Database | Messier, J., B. J. McGill, and M. J. Lechowicz. 2010. How do traits vary across ecological scales? A case for trait-based ecology. Ecology Letters 13:838-848. |
| Panama Plant Traits Database | Wright, S. J., K. Kitajima, N. J. B. Kraft, P. B. Reich, I. J. Wright, D. E. Bunker, R. Condit, J. W. Dalling, S. J. Davies, S. Díaz, B. M. J. Engelbrecht, K. E. Harms, S. P. Hubbell, C. O. Marks, M. C. Ruiz-Jaen, C. M. Salvador, and A. E. Zanne. 2011 . Functional traits and the growth-mortality tradeoff in tropical trees. Ecology 91:3664-3674. |
| Panama Tree Traits | Craven, D., D. Braden, M. S. Ashton, G. P. Berlyn, M. Wishnie, and D. Dent. 2007. Between and within-site comparisons of structural and physiological characteristics and foliar nutrient content of 14 tree species at a wet, fertile site and a dry, infertile site in Panama. Forest Ecology and Management 238:335-346. |
| Photosynthetic Capacity Dataset | Carswell, F. E., Meir, P., Wandelli, E. V., Bonates, L. C. M., Kruijt, B., Barbosa, E. M., Nobre, A. D. & Jarvis, P. G. 2000 Photosynthetic capacity in a central Amazonian rain forest. Tree physiology. 20, 3, p. 179-186 8 p. |
| PLANTSdata USDA | Green, W. 2009. USDA PLANTS Compilation, version 1, 09-02-02. (http://bricol.net/downloads/data/PLANTSdatabase/) NRCS: The PLANTS Database (http://plants.usda.gov, 1 Feb 2009). National Plant Data Center: Baton Rouge, LA 70874-74490 USA. |
| The Functional Ecology of Trees (FET) Database - Jena | Wirth, C. and J. W. Lichstein. 2009. The Imprint of Species Turnover on Old-Growth Forest Carbon Balances - Insights From a Trait-Based Model of Forest Dynamics. Pages 81-113 in C. Wirth, G. Gleixner, and M. Heimann, editors. Old-Growth Forests: Function, Fate and Value. Springer, New York, Berlin, Heidelberg. |
| The Tansley Review LMA Database | Poorter, H., U. Niinemets, L. Poorter, I. J. Wright, and R. Villar. 2009. Causes and consequences of variation in leaf mass per area (LMA): a meta-analysis. New Phytologist 182:565-588. |
| The Xylem/Phloem Database | Schweingruber, F.H., Landolt, W.: The Xylem Database. Swiss Federal Research Institute WSL Updated (2005) |
| Xylem Functional Traits (XFT) Database | Brendan Choat, Steven Jansen, Tim J. Brodribb, Herve Cochard, Sylvain Delzon, Radika Bhaskar, Sandra J. Bucci, Taylor S. Feild, Sean M. Gleason, Uwe G. Hacke, Anna L. Jacobsen, Frederic Lens, Hafiz Maherali, Jordi Martinez-Vilalta, Stefan Mayr, Maurizio Mencuccini, Patrick J. Mitchell, Andrea Nardini, Jarmila Pittermann, R. Brandon Pratt, John S. Sperry, Mark Westoby, Ian J. Wright & Amy E. Zanne (2012) Global convergence in the vulnerability of forests to drought. Nature 491:752-755 doi:10.1038/nature11688 |
| Yasuni Ecuador Leaves | Kraft, N. J. B., R. Valencia, and D. Ackerly. 2008. Functional traits and niche-based tree community assembly in an Amazonian forest. Science 322:580-582. |

**APPENDIX 2.** List of the books from which additional functional trait information was obtained.

Carvalho, P. E. R. (2003). Espécies Arbóreas Brasileiras- Volume 1 (1ª ed.). Brasília-DF. Embrapa.

Carvalho, P. E. R. (2006). Espécies Arbóreas Brasileiras- Volume 2 (1ª ed.). Brasília-DF. Embrapa.

Carvalho, P. E. R. (2008). Espécies Arbóreas Brasileiras- Volume 3 (1ª ed.). Brasília-DF. Embrapa.

Carvalho, P. E. R. (2010). Espécies Arbóreas Brasileiras- Volume 4 (1ª ed.). Brasília-DF. Embrapa.

Carvalho, P. E. R. (2014). Espécies Arbóreas Brasileiras- Volume 5 (1ª ed.). Brasília-DF. Embrapa.

Kuhlmann, M. (2012). Frutos e Sementes do cerrado - Atrativos para a Fauna - Guia de campo. Brasília-DF. Rede de sementes do cerrado.

Lorenzi, H. (2000). Árvores Brasileiras - Manual de Identificação e Cultivo de Plantas Nativas do Brasil- Vol.1 (3ª ed.). Nova Odessa-SP. Instituto Plantarum de Estudos da Flora LTDA.

Lorenzi, H. (2000). Árvores Brasileiras - Manual de Identificação e Cultivo de Plantas Nativas do Brasil- Vol.2 (3ª ed.). Nova Odessa-SP. Instituto Plantarum de Estudos da Flora LTDA.

Lorenzi, H. (2000). Árvores Brasileiras - Manual de Identificação e Cultivo de Plantas Nativas do Brasil- Vol.3 (3ª ed.). Nova Odessa-SP. Instituto Plantarum de Estudos da Flora LTDA.

Silva Júnior, M. C. (2005). 100 árvores do cerrado. Brasília-DF. Rede de sementes do cerrado.

Silva Júnior, M. C..; Pereira, B. A. S. (2009). Mais 100 árvores do cerrado matas de galeria - Guia de campo. Brasília-DF. Rede de sementes do cerrado.

Silva Júnior, M. C.; Lima. R. M. C. (2010). 100 árvores urbanas Brasília - Guia de campo. Brasília-DF. Rede de sementes do cerrado.

**APPENDIX 3.** List of the references used to obtain data from Cerrado woody riparian communities.

| **Site** | **Reference** |
| --- | --- |
| Corrego Bacaba Alto | Miguel, A., Marimon, B. S., Oliveira, E. A. de, Maracahipes, L., & Marimon-Junior, B. H. (2011). Dinâmica da comunidade lenhosa de uma floresta de galeria na transição Cerrado-Floresta Amazônica no Leste de Mato Grosso, em um período de sete anos (1999 a 2006). Biota Neotropica, 11(1), 53–61. doi:10.1590/S1676-06032011000100005 |
| Corrego Bacaba Meio | Miguel, A., Marimon, B. S., Oliveira, E. A. de, Maracahipes, L., & Marimon-Junior, B. H. (2011). Dinâmica da comunidade lenhosa de uma floresta de galeria na transição Cerrado-Floresta Amazônica no Leste de Mato Grosso, em um período de sete anos (1999 a 2006). Biota Neotropica, 11(1), 53–61. doi:10.1590/S1676-06032011000100005 |
| EPA-Pandeiros Balneario | de Azevedo, I. F. P., Nunes, Y. R. F., de Ávila, M. A., da Silva, D. L., Fernandes, G. W., & Veloso, R. B. (2014). Phenology of riparian tree species in a transitional region in southeastern Brazil. Revista Brasileira de Botanica, 37(1), 47–59. doi:10.1007/s40415-014-0046-5 |
| EPA-Pandeiros Faz.Agropop | de Azevedo, I. F. P., Nunes, Y. R. F., de Ávila, M. A., da Silva, D. L., Fernandes, G. W., & Veloso, R. B. (2014). Phenology of riparian tree species in a transitional region in southeastern Brazil. Revista Brasileira de Botanica, 37(1), 47–59. doi:10.1007/s40415-014-0046-5 |
| EPA-Pandeiros Sao Domingos | de Azevedo, I. F. P., Nunes, Y. R. F., de Ávila, M. A., da Silva, D. L., Fernandes, G. W., & Veloso, R. B. (2014). Phenology of riparian tree species in a transitional region in southeastern Brazil. Revista Brasileira de Botanica, 37(1), 47–59. doi:10.1007/s40415-014-0046-5 |
| Fazenda Agua Limpa | Ribeiro, G. H. P. M., & Felfili, J. M. (2009) . Regeneração Natural Em Diferentes Ambientes Da Mata de Galeria Do Capetinga, Na Fazenda Água Limpa-DF, Cerne, 15(1), 1–9. |
| IBGE | Paiva, A. O., Silva, L. C. R., & Haridasan, M. (2015). Productivity-efficiency tradeoffs in tropical gallery forest-savanna transitions: linking plant and soil processes through litter input and composition. Plant Ecology, 216(6), 775–787. doi:10.1007/s11258-015-0466-8 |
| Parque da Matinha | Prado Júnior, J. A., Lopes, S. D. F., Vale, V. S., Dias Neto, O. C., & Schiavini, I. (2012). Comparação florística, estrutural e ecológica da vegetação arbórea das fitofisionomias de um remanescente urbano de cerrado. Biosceince Journal, 24(3) 456–471. |
| PNSC Brejo Velho | Matos, M. D. Q., & Felfili, J. M. (2010). Florística, fitossociologia e diversidade da vegetação arbórea nas matas de galeria do Parque Nacional de Sete Cidades (PNSC), Piauí, Brasil. Acta Botanica Brasilica, 24(2), 483–496. doi:10.1590/S0102-33062010000200019 |
| PNSC Mata da Sambaiba | Matos, M. D. Q., & Felfili, J. M. (2010). Florística, fitossociologia e diversidade da vegetação arbórea nas matas de galeria do Parque Nacional de Sete Cidades (PNSC), Piauí, Brasil. Acta Botanica Brasilica, 24(2), 483–496. doi:10.1590/S0102-33062010000200019 |
| PNSC Mata do Bacuri | Matos, M. D. Q., & Felfili, J. M. (2010). Florística, fitossociologia e diversidade da vegetação arbórea nas matas de galeria do Parque Nacional de Sete Cidades (PNSC), Piauí, Brasil. Acta Botanica Brasilica, 24(2), 483–496. doi:10.1590/S0102-33062010000200019 |
| Rio Cuiaba P1 | Umetsu, R. K., Girard, P., Matos, D. M. da S., & Silva, C. J. da. (2011). Efeito da inundação lateral sobre a distribuição da vegetação ripária em um trecho do rio Cuiabá, MT. Revista Árvore, 35(5), 1077–1087. doi:10.1590/S0100-67622011000600014 |
| Rio Cuiaba P2 | Umetsu, R. K., Girard, P., Matos, D. M. da S., & Silva, C. J. da. (2011). Efeito da inundação lateral sobre a distribuição da vegetação ripária em um trecho do rio Cuiabá, MT. Revista Árvore, 35(5), 1077–1087. doi:10.1590/S0100-67622011000600014 |
| Rio Cuiaba P3 | Umetsu, R. K., Girard, P., Matos, D. M. da S., & Silva, C. J. da. (2011). Efeito da inundação lateral sobre a distribuição da vegetação ripária em um trecho do rio Cuiabá, MT. Revista Árvore, 35(5), 1077–1087. doi:10.1590/S0100-67622011000600014 |
| Rio Cuiaba P4 | Umetsu, R. K., Girard, P., Matos, D. M. da S., & Silva, C. J. da. (2011). Efeito da inundação lateral sobre a distribuição da vegetação ripária em um trecho do rio Cuiabá, MT. Revista Árvore, 35(5), 1077–1087. doi:10.1590/S0100-67622011000600014 |
| Rio Cuiaba P5 | Umetsu, R. K., Girard, P., Matos, D. M. da S., & Silva, C. J. da. (2011). Efeito da inundação lateral sobre a distribuição da vegetação ripária em um trecho do rio Cuiabá, MT. Revista Árvore, 35(5), 1077–1087. doi:10.1590/S0100-67622011000600014 |
| Rio Cuiaba P9 | Umetsu, R. K., Girard, P., Matos, D. M. da S., & Silva, C. J. da. (2011). Efeito da inundação lateral sobre a distribuição da vegetação ripária em um trecho do rio Cuiabá, MT. Revista Árvore, 35(5), 1077–1087. doi:10.1590/S0100-67622011000600014 |
| Rio Guariroba | de Oliveira, A. K. M., Resende, U. M., & Ribeiro, F. D. (2006). Estrutura Arbórea De Um Trecho De Mata Ciliar No Município De Campo Grande, MS. Ensaios E Ciência, 10(1), 133-141. |
| Vale do Veu de Noiva 1 | Pinto, J. R. R., & Oliveira-Filho, A. T. (2005). Mudanças florísticas e estruturais na comunidade arbórea de uma floresta de vale no Parque Nacional da Chapada dos Guimarães, Mato Grosso, Brasil. Revista Brasileira de Botânica, 28(3), 523–539. doi:10.1590/S0100-84042005000300010 |
| Vale do Veu de Noiva 2 | Pinto, J. R. R., & Oliveira-Filho, A. T. (2005). Mudanças florísticas e estruturais na comunidade arbórea de uma floresta de vale no Parque Nacional da Chapada dos Guimarães, Mato Grosso, Brasil. Revista Brasileira de Botânica, 28(3), 523–539. doi:10.1590/S0100-84042005000300010 |

**APPENDIX 4.** List of species from the 24 study communities.

*Aegiphila brachiata, Aegiphila verticillata, Agonandra brasiliensis, Albizia niopoides, Alchornea castaneifolia, Alchornea discolor, Alchornea glandulosa, Alibertia edulis, Allophylus edulis, Allophylus semidentatus, Amaioua guianensis, Anacardium occidentale, Anadenanthera colubrina, Andira vermifuga, Annona montana, Apeiba tibourbou, Apuleia leiocarpa, Aspidosperma australe, Aspidosperma cylindrocarpon, Aspidosperma multiflorum, Aspidosperma spruceanum, Aspidosperma subincanum, Aspidosperma tomentosum, Astrocaryum aculeatum, Astrocaryum vulgare, Astronium fraxinifolium, Astronium graveolens, Attalea phalerata, Averrhoidium gardnerianum, Banara tomentosa, Bauhinia longifolia, Bauhinia rufa, Bellucia grossularioides, Blepharocalyx salicifolius, Bocageopsis mattogrossensis, Brosimum gaudichaudii, Buchenavia tetraphylla, Byrsonima crassifolia, Byrsonima laxiflora, Byrsonima pachyphylla, Byrsonima sericea, Byrsonima verbascifolia, Callisthene fasciculata, Callisthene major, Calophyllum brasiliense, Calyptranthes strigipes, Campomanesia aromatica, Campomanesia guazumifolia, Campomanesia velutina, Cardiopetalum calophyllum, Cariniana estrellensis, Caryocar brasiliense, Casearia arborea, Casearia gossypiosperma, Casearia lasiophylla, Casearia sylvestris, Cecropia pachystachya, Cedrela fissilis, Ceiba speciosa, Cheiloclinium cognatum, Chloroleucon tenuiflorum, Chloroleucon tortum, Chrysophyllum amazonicum, Chrysophyllum gonocarpum, Chrysophyllum marginatum, Coccoloba mollis, Combretum leprosum, Connarus perrottetii, Connarus suberosus, Copaifera coriacea, Copaifera langsdorffii, Cordia bicolor, Cordia naidophila, Cordia sellowiana, Cordia trichotoma, Cordiera macrophylla, Couepia grandiflora, Coussarea hydrangeifolia, Cryptocarya aschersoniana, Cupania vernalis, Curatella americana, Cybianthus brasiliensis, Cybianthus cuneifolius, Cybianthus glaber, Cybianthus guyanensis, Dalbergia foliolosa, Dalbergia miscolobium, Davilla elliptica, Dendropanax cuneatus, Dilodendron bipinnatum, Dimorphandra gardneriana, Dimorphandra mollis, Diospyros guianensis, Diospyros hispida, Diospyros sericea, Dipteryx alata, Diptychandra aurantiaca, Duguetia echinophora, Elaeoluma glabrescens, Endlicheria paniculata, Enterolobium gummiferum, Ephedranthus parviflorus, Ephedranthus pisocarpus, Eremanthus glomerulatus, Eriotheca gracilipes, Eriotheca pubescens, Erythroxylum anguifugum, Erythroxylum daphnites, Erythroxylum suberosum, Erythroxylum tortuosum, Eugenia florida, Eugenia uniflora, Euplassa inaequalis, Euterpe precatoria, Faramea hyacinthina, Ficus enormis, Ficus gomelleira, Ficus guaranitica, Ficus insipida, Ficus krukovii, Ficus maxima, Ficus pertusa, Gallesia integrifolia, Garcinia brasiliensis, Guapira areolata, Guapira noxia, Guapira opposita, Guarea guidonia, Guarea kunthiana, Guarea macrophylla, Guatteria sellowiana, Guazuma ulmifolia, Guettarda viburnoides, Handroanthus impetiginosus, Handroanthus ochraceus, Handroanthus serratifolius, Heisteria densifrons, Heisteria ovata, Himatanthus bracteatus, Hirtella glandulosa, Hirtella gracilipes, Hyeronima alchorneoides, Hymenaea courbaril, Hymenaea eriogyne, Hymenaea stigonocarpa, Inga alba, Inga cayennensis, Inga cylindrica, Inga edulis, Inga laurina, Inga marginata, Inga vera, Ixora brevifolia, Jacaranda copaia, Jacaranda cuspidifolia, Jacaranda puberula, Kielmeyera coriacea, Leptolobium dasycarpum, Licania apetala, Licania blackii, Licania hoehnei, Licania kunthiana, Licania parvifolia, Licania sclerophylla, Lithrea molleoides, Lonchocarpus cultratus, Luehea candicans, Luehea paniculata, Mabea fistulifera, Mabea pohliana, Machaerium acutifolium, Machaerium brasiliense, Machaerium opacum, Machaerium villosum, Maclura tinctoria, Magonia pubescens, Manihot tripartita, Maprounea guianensis, Margaritaria nobilis, Matayba elaeagnoides, Matayba guianensis, Maytenus floribunda, Mezilaurus crassiramea, Miconia albicans, Miconia chartacea, Miconia elegans, Miconia ferruginata, Miconia longifolia, Miconia matthaei, Miconia minutiflora, Miconia nervosa, Miconia punctata, Miconia splendens, Micropholis venulosa, Mimosa adenocarpa, Mollinedia schottiana, Mouriri glazioviana, Mouriri guianensis, Mouriri pusa, Myracrodruon urundeuva, Myrcia fenzliana, Myrcia hebepetala, Myrcia multiflora, Myrcia splendens, Myrcia tomentosa, Myrciaria floribunda, Myrsine coriacea, Myrsine guianensis, Myrsine lancifolia, Myrsine umbellata, Nectandra cissiflora, Nectandra cuspidata, Nectandra nitidula, Neea theifera, Ocotea aciphylla, Ocotea corymbosa, Ocotea elegans, Ocotea minarum, Ocotea pomaderroides, Ocotea pulchella, Ocotea spixiana, Ocotea velloziana, Oenocarpus distichus, Ormosia arborea, Ormosia fastigiata, Ormosia stipularis, Ouratea hexasperma, Oxandra sessiliflora, Palicourea rigida, Parkia platycephala, Pera glabrata, Persea fusca, Piper amalago, Piper arboreum, Piper tuberculatum, Piptadenia gonoacantha, Piptocarpha macropoda, Plathymenia reticulata, Platypodium elegans, Plenckia populnea, Pleradenophora membranifolia, Plinia peruviana, Poecilanthe parviflora, Pouteria gardneri, Pouteria glomerata, Pouteria ramiflora, Pouteria torta, Priogymnanthus hasslerianus, Protium guianense, Protium heptaphyllum, Protium spruceanum, Pseudobombax longiflorum, Pseudobombax tomentosum, Pseudolmedia laevigata, Psidium myrtoides, Psidium salutare, Pterodon pubescens, Qualea dichotoma, Qualea grandiflora, Qualea multiflora, Qualea parviflora, Randia armata, Rhamnidium elaeocarpum, Richeria grandis, Roupala montana, Rudgea viburnoides, Sapium glandulosum, Schefflera macrocarpa, Schefflera morototoni, Senegalia polyphylla, Simarouba amara, Simarouba versicolor, Siparuna guianensis, Siphoneugena densiflora, Sorocea guilleminiana, Sparattosperma leucanthum, Spondias mombin, Sterculia apetala, Sterculia striata, Stryphnodendron adstringens, Stryphnodendron coriaceum, Styrax camporum, Styrax ferrugineus, Styrax pohlii, Symphonia globulifera, Tabebuia aurea, Tabebuia roseoalba, Tachigali paniculata, Tachigali vulgaris, Talisia esculenta, Tapirira guianensis, Tapirira obtusa, Tapura amazonica, Terminalia argentea, Terminalia fagifolia, Terminalia glabrescens, Terminalia phaeocarpa, Tetragastris altissima, Tetragastris cerradicola, Tibouchina candolleana, Tocoyena brasiliensis, Tocoyena formosa, Trichilia clausseni, Trichilia elegans, Trichilia pallida, Triplaris americana, Unonopsis guatterioides, Urera baccifera, Vatairea macrocarpa, Virola albidiflora, Virola sebifera, Virola surinamensis, Vochysia divergens, Vochysia haenkeana, Vochysia tucanorum, Ximenia americana, Xylopia aromatica, Xylopia emarginata.*

**Appendix 5.** Maximum Functional Richness (FRic_max_) by species proportion of each solution in each evaluated riparian woody community.

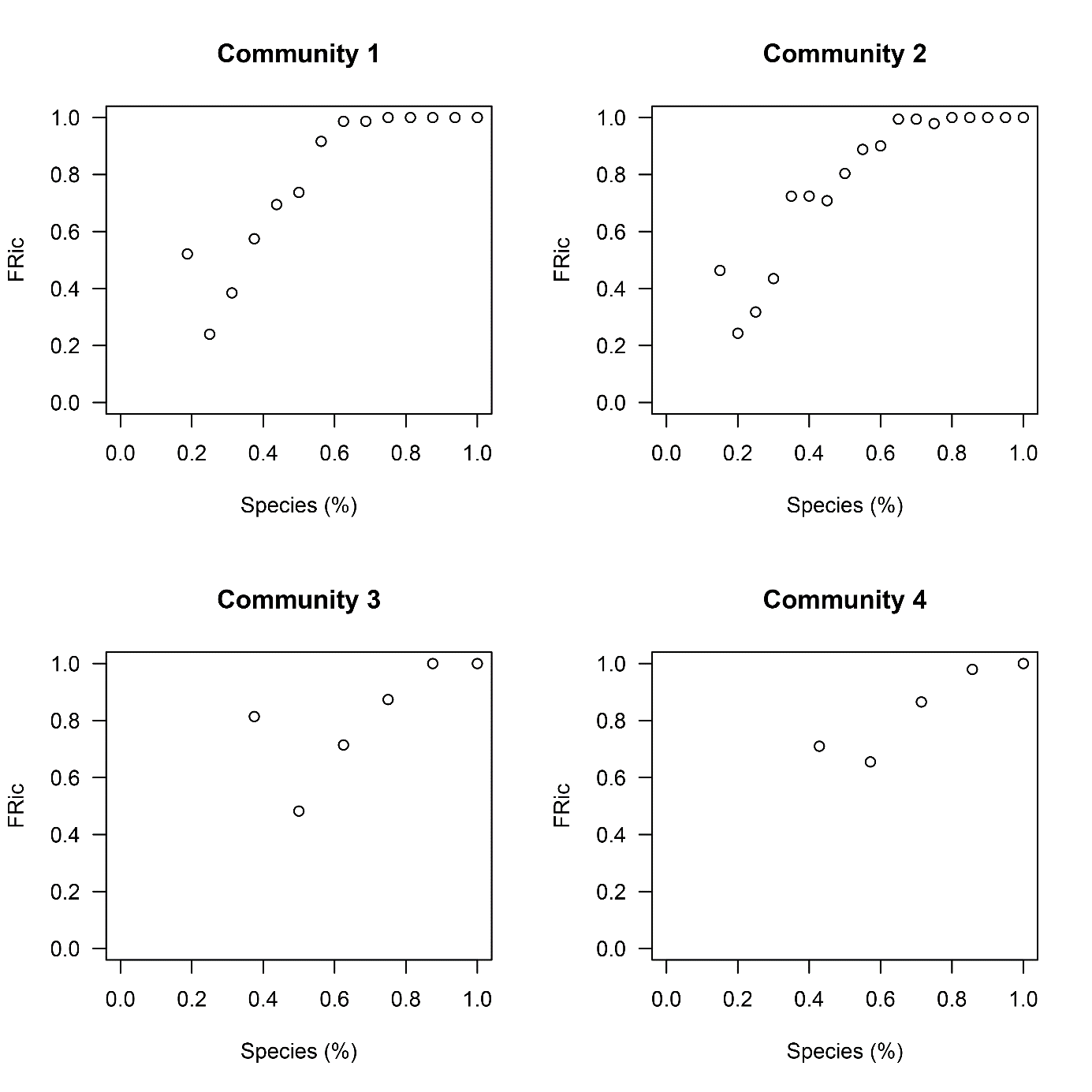

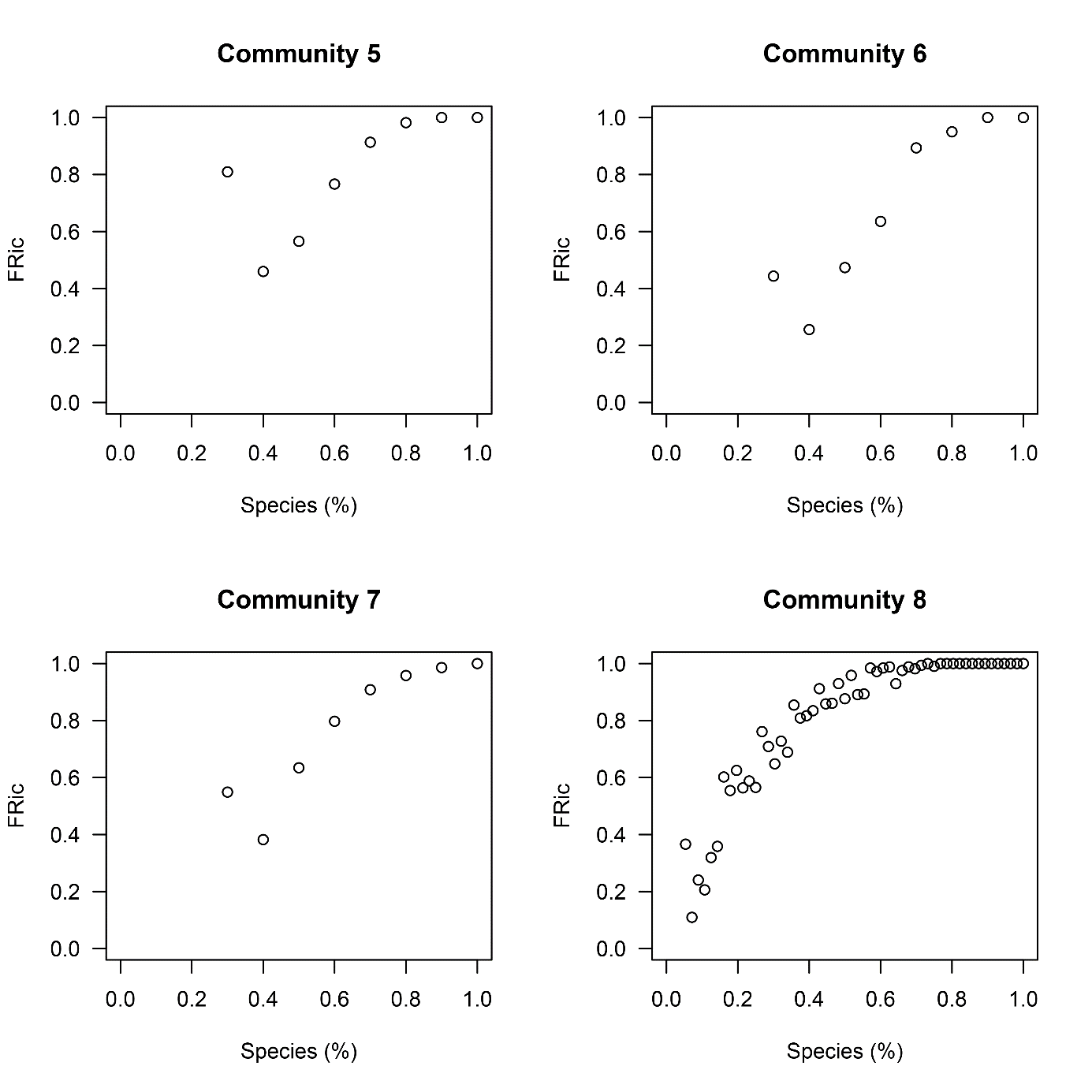

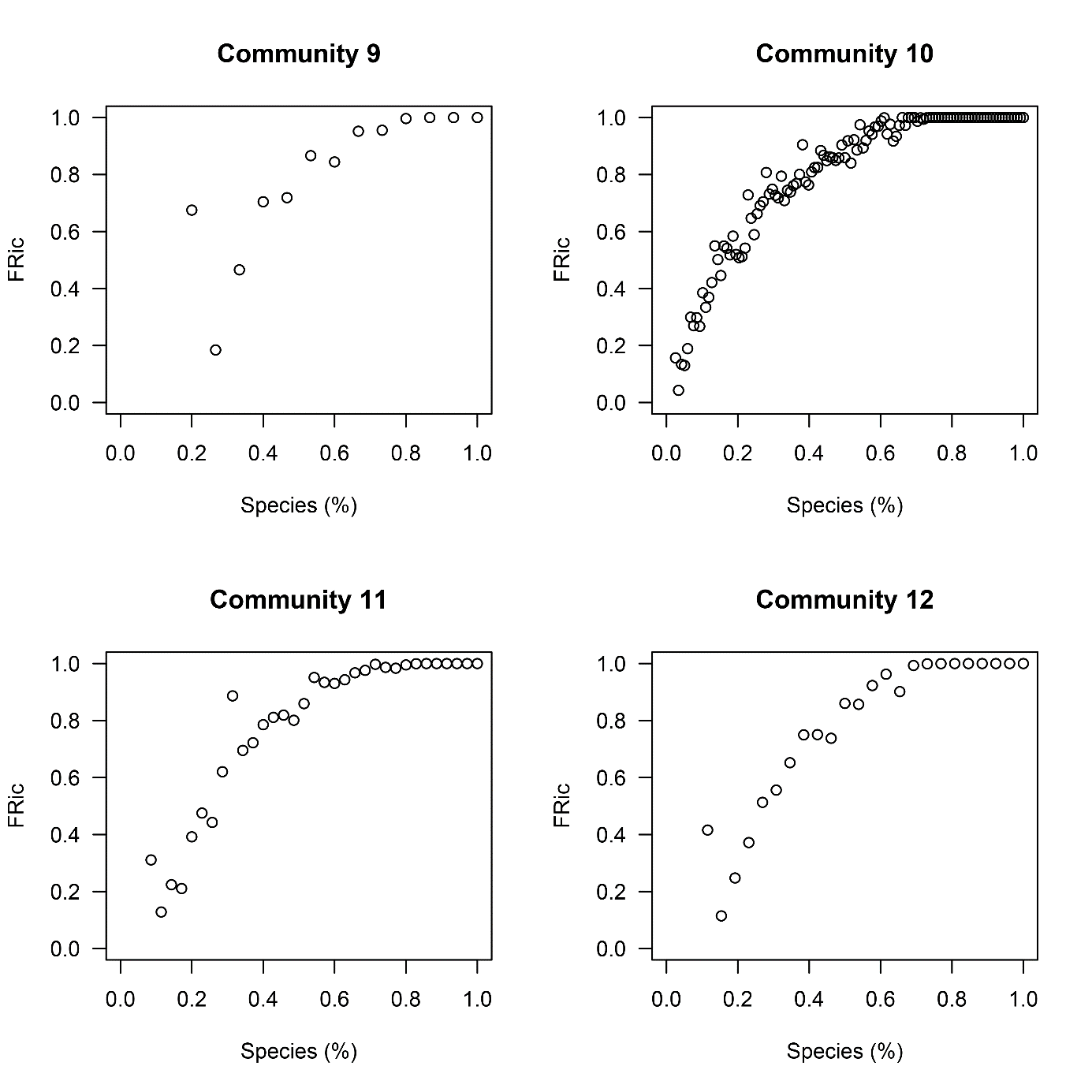

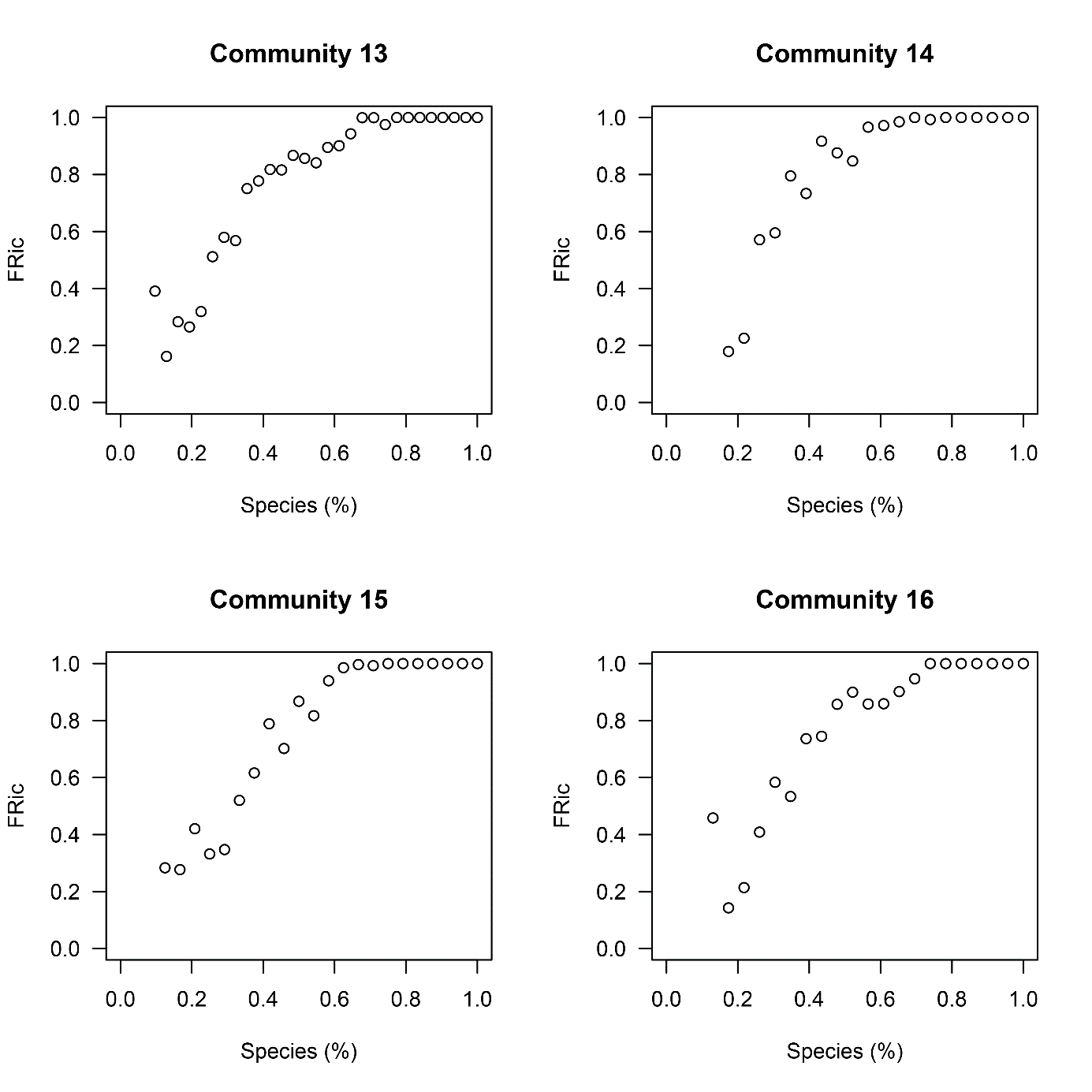

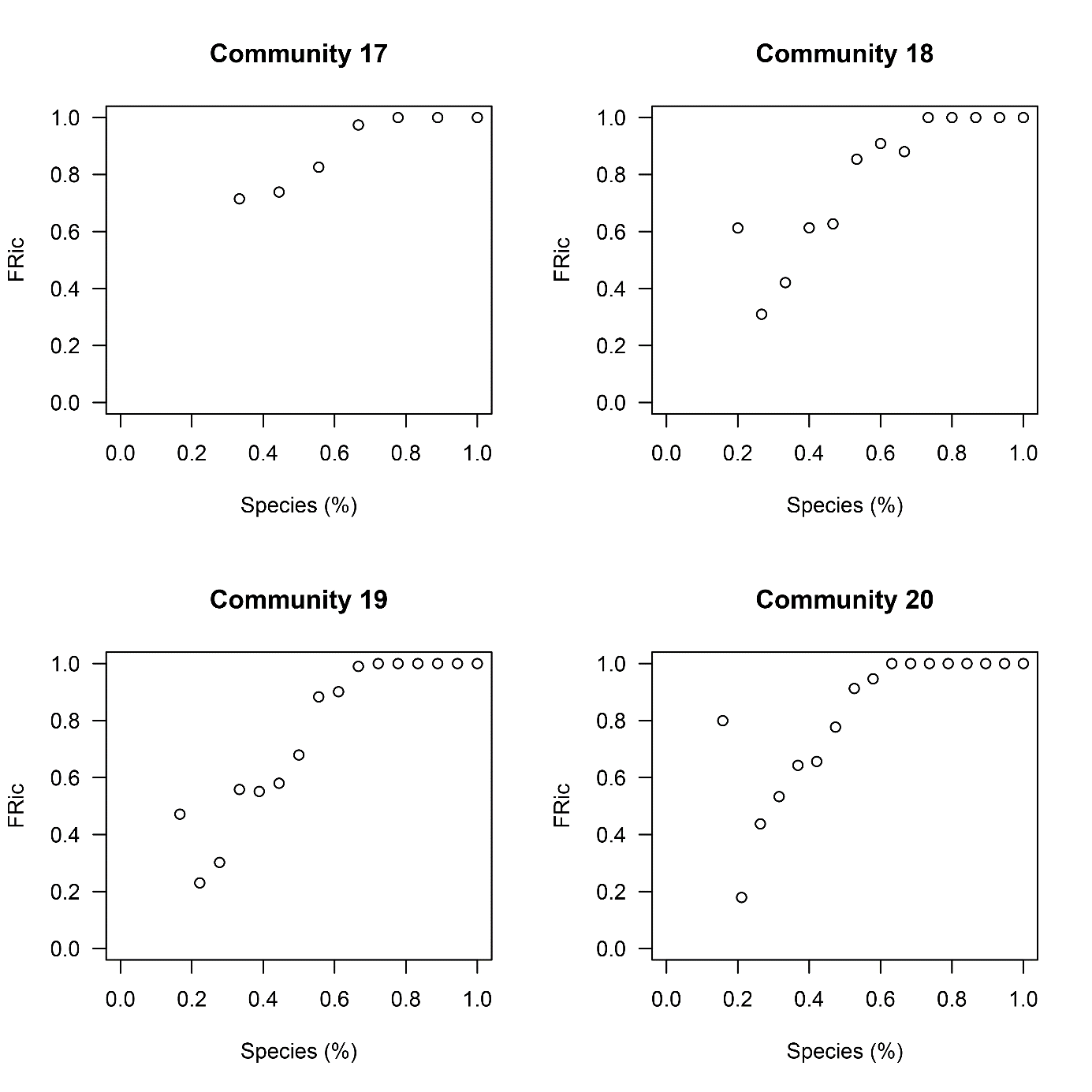

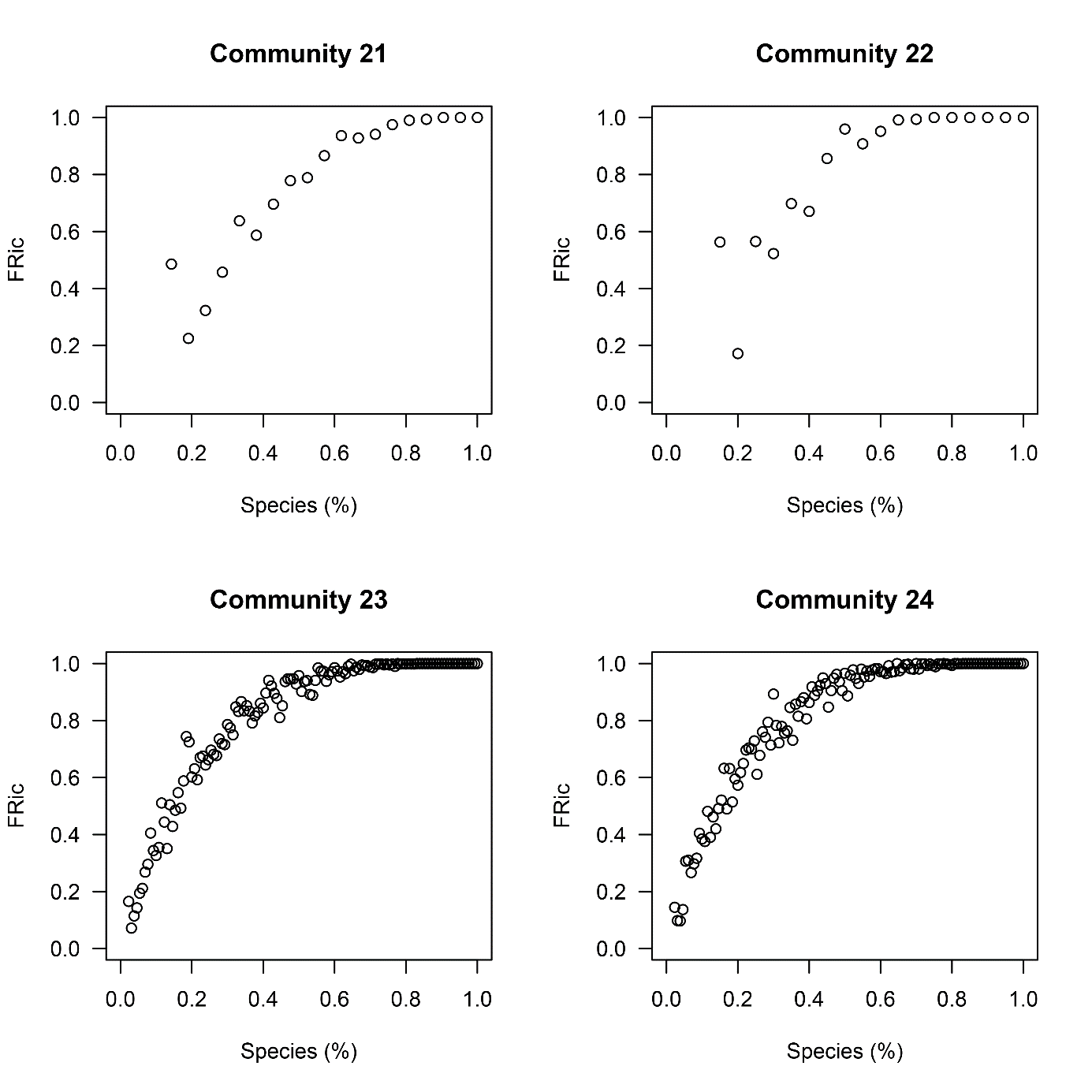

**Appendix 6.** Maximum Functional Redundancy (FR_max_) by species proportion of each solution in each evaluated riparian woody community.

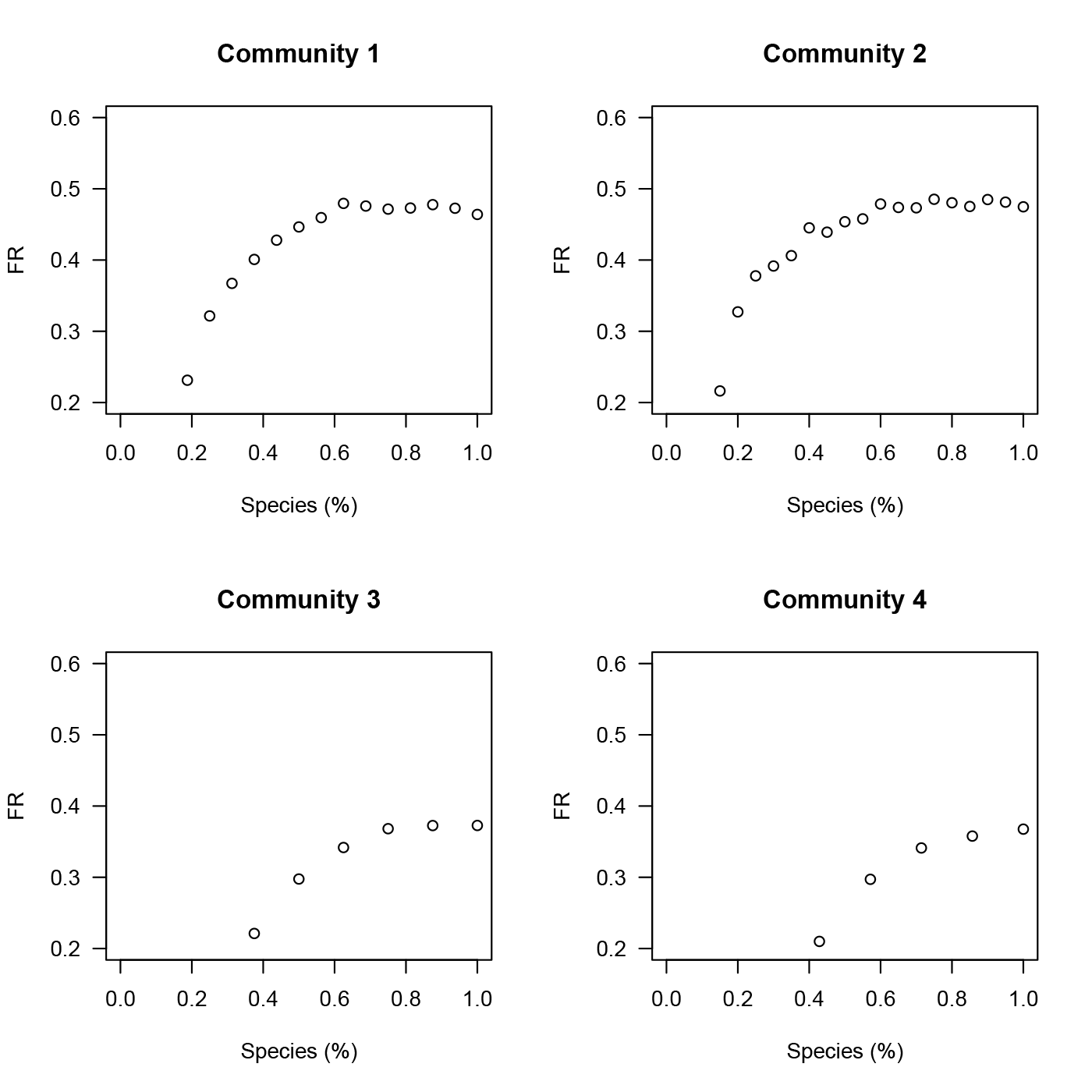

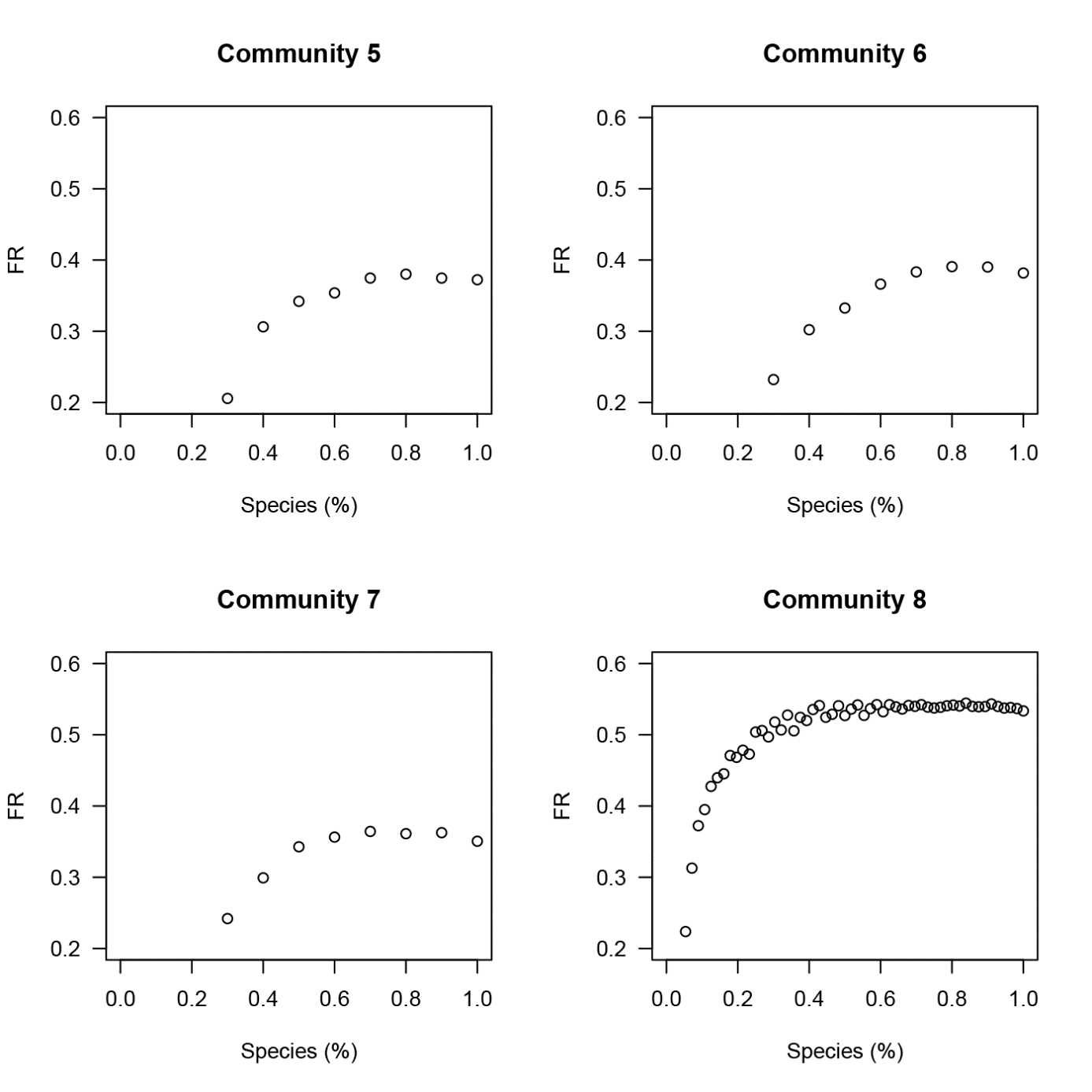

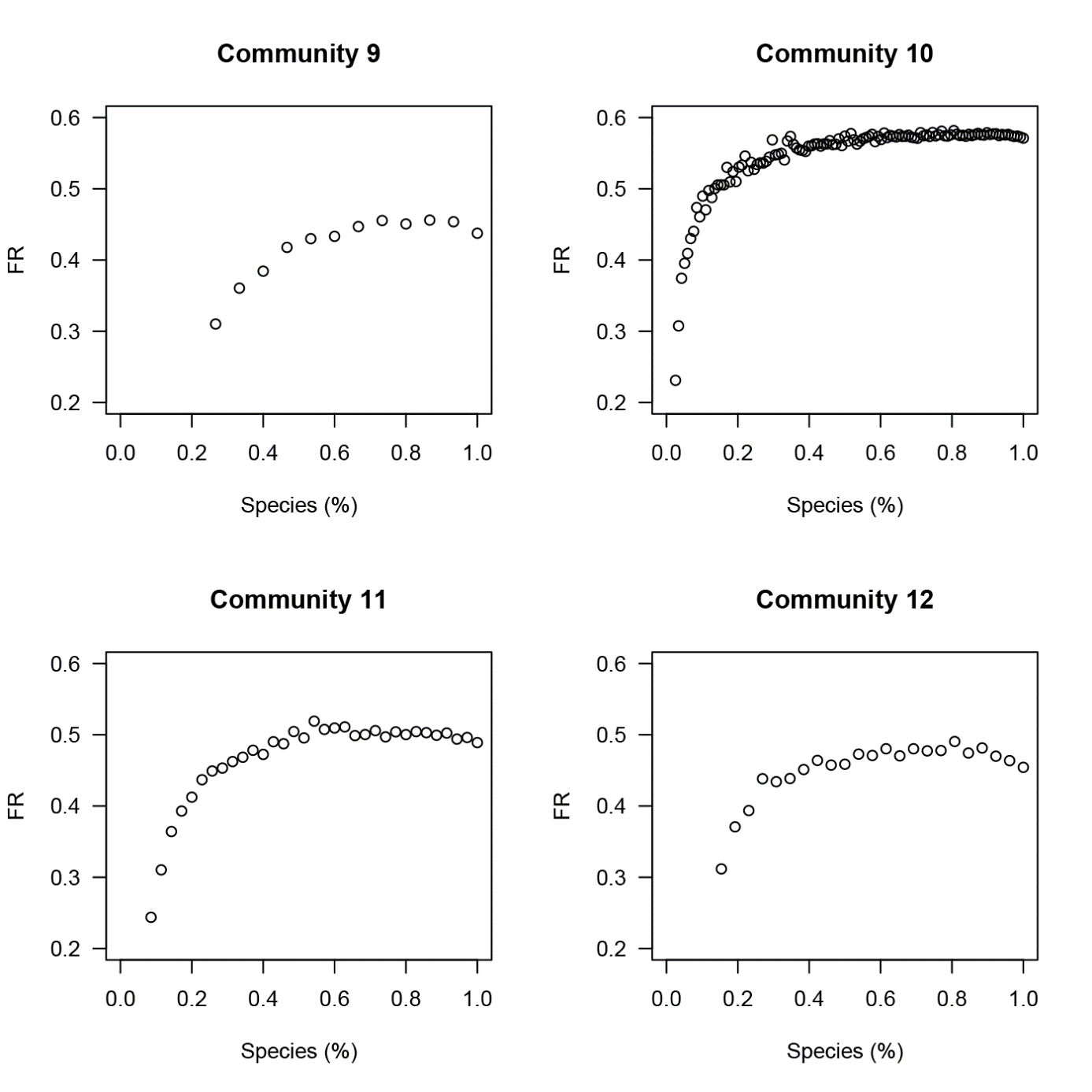

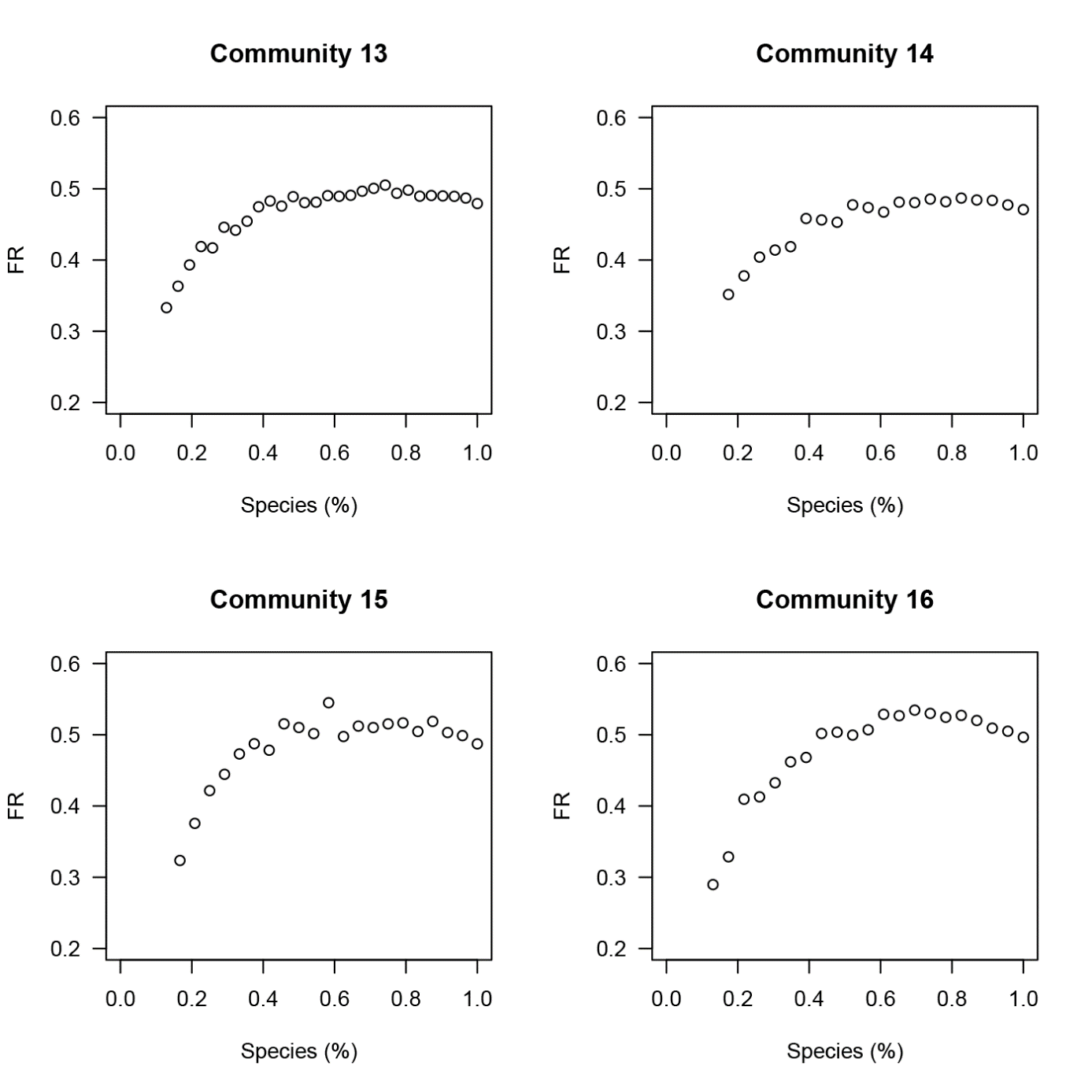

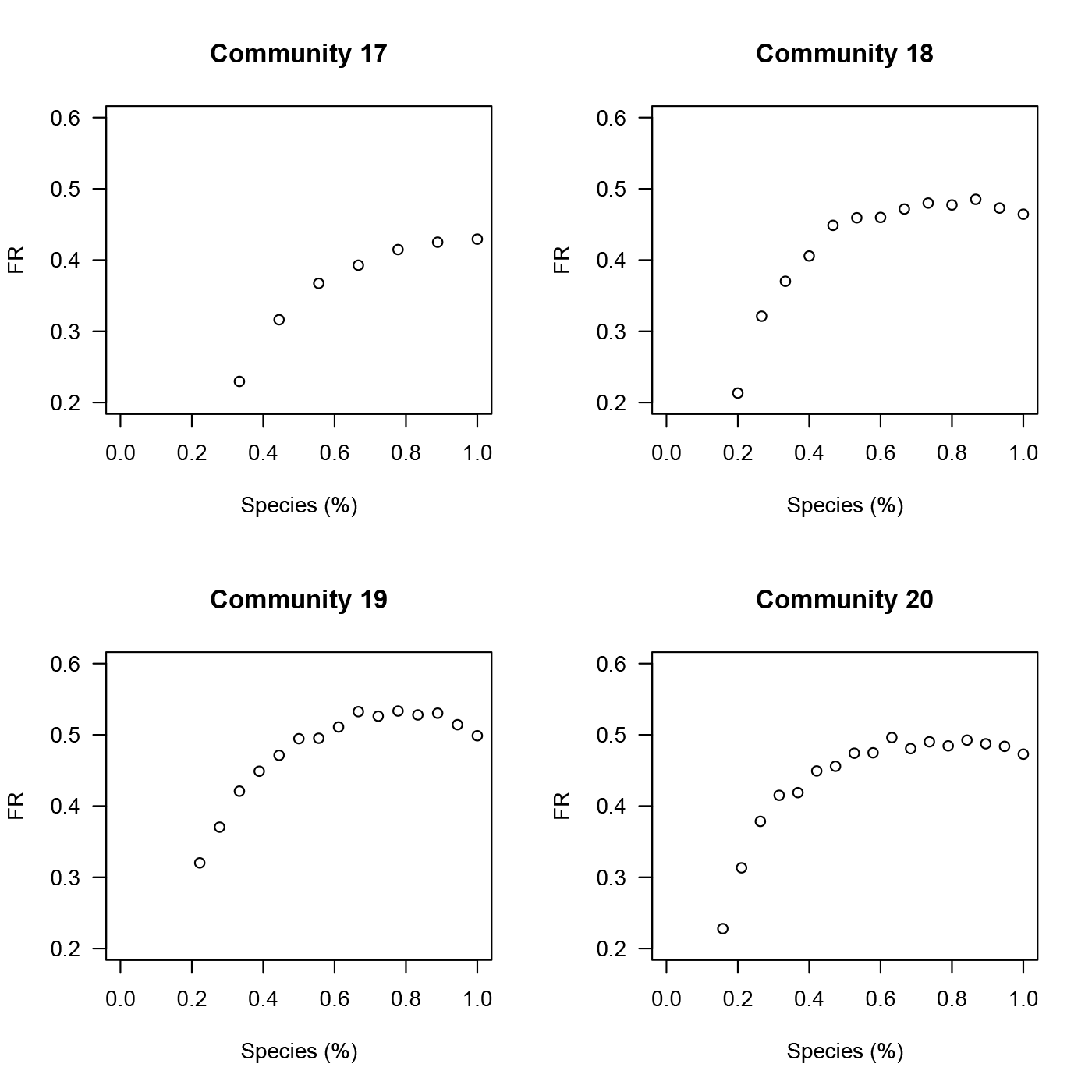

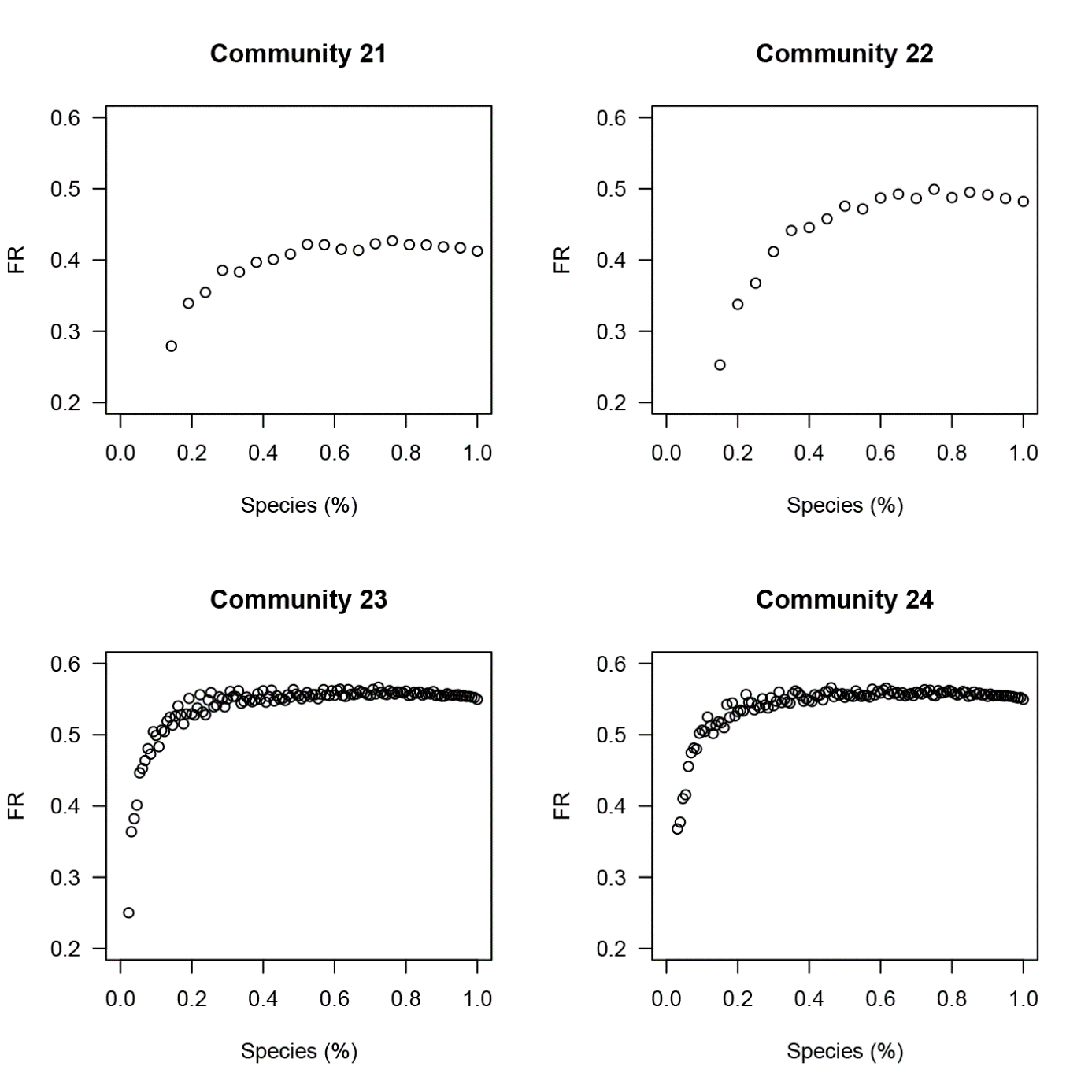

**Appendix 8 –** Comparison between FRic in two scenarios: calculated with the set of species of community that maximizes FRic, and with the set of species that maximizes FR. Points represent FRic of the 24 communities (*W* = 554, *p* < 0.001).

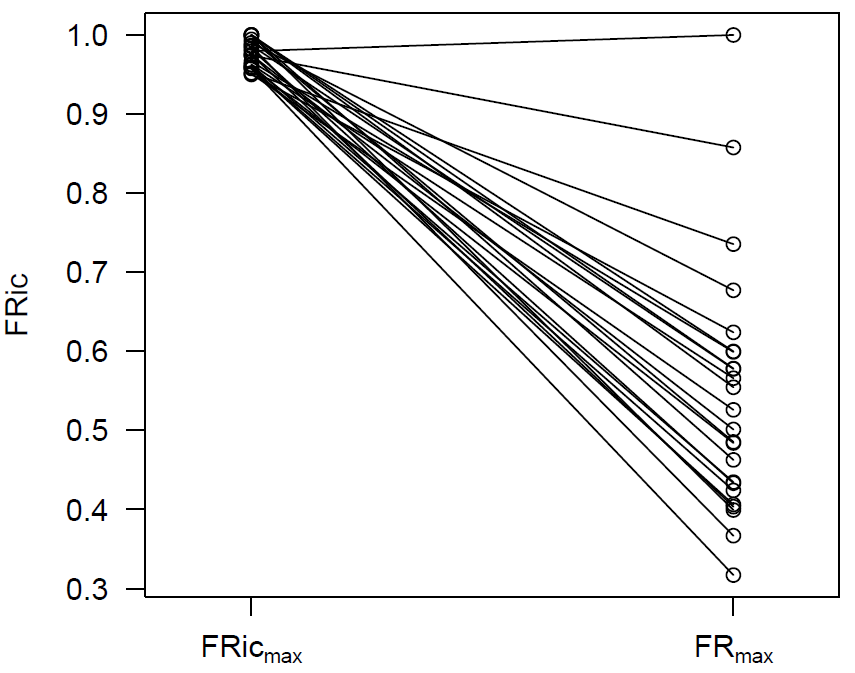
